# Human Endometrial Transcriptome and Progesterone Receptor Cistrome Reveal Important Pathways and Epithelial Regulators

**DOI:** 10.1101/680181

**Authors:** Ru-pin Alicia Chi, Tianyuan Wang, Nyssa Adams, San-pin Wu, Steven L. Young, Thomas E. Spencer, Francesco DeMayo

## Abstract

**Context:** Poor uterine receptivity is one major factor leading to pregnancy loss and infertility. Understanding the molecular events governing successful implantation is hence critical in combating infertility.

**Objective:** To define PGR-regulated molecular mechanisms and epithelial roles in receptivity.

**Design:** RNA-seq and PGR-ChIP-seq were conducted in parallel to identify PGR-regulated pathways during the WOI in endometrium of fertile women.

**Setting:** Endometrial biopsies from the proliferative and mid-secretory phases were analyzed.

**Patients or Other Participants:** Participants were fertile, reproductive aged (18-37) women with normal cycle length; and without any history of dysmenorrhea, infertility, or irregular cycles. In total, 42 endometrial biopsies obtained from 42 women were analyzed in this study.

**Interventions:** There were no interventions during this study.

**Main Outcome Measures:** Here we measured the alterations in gene expression and PGR occupancy in the genome during the WOI, based on the hypothesis that PGR binds uterine chromatin cycle-dependently to regulate genes involved in uterine cell differentiation and function.

**Results:** 653 genes were identified with regulated PGR binding and differential expression during the WOI. These were involved in regulating inflammatory response, xenobiotic metabolism, EMT, cell death, interleukin/STAT signaling, estrogen response, and MTORC1 response. Transcriptome of the epithelium identified 3,052 DEGs, of which 658 were uniquely regulated. Transcription factors IRF8 and MEF2C were found to be regulated in the epithelium during the WOI at the protein level, suggesting potentially important functions that are previously unrecognized.

**Conclusion:** PGR binds the genomic regions of genes regulating critical processes in uterine receptivity and function.

**Précis:** Using a combination of RNA-seq and PGR ChIP-seq, novel signaling pathways and epithelial regulators were identified in the endometrium of fertile women during the window of implantation.

## Introduction

The human endometrium is a highly complex tissue. The functionalis layer consists of the stromal compartment which makes up significant portion of the endometrium; the glandular epithelium which is responsible for secreting an array of growth factors and cytokines (1); and the luminal epithelium which lines the stromal compartment and is the first maternal interface with which the embryo interacts inside the uterus. In order to maximize the chances of a successful pregnancy, the uterus prepares for embryo implantation after each menstruation by the generation and differentiation of the endometrial functionalis, a process known as the menstrual cycle (2, 3). This is orchestrated by the interplay of two steroid hormones, estrogen and progesterone. During the proliferative (P) phase, estrogen promotes proliferation of both the stromal and epithelial cells, steadily increasing the thickness of the functionalis (4, 5). Upon ovulation, the ovary begins secreting progesterone, halting estrogen-induced proliferation and initiating differentiation of stromal cells (decidualization) and epithelial cells. These include depolarization, altered surface morphology, expression of specific adhesion proteins, altered steroid receptor expression, and secretion of glycogen (5, 6). Without a successful implantation, the levels of both steroid hormones decrease during the late secretory phase, leading to endometrial involution and subsequently endometrial shedding (menstruation), initiating another cycle (7).

Abnormal embryo implantation and implantation failure are major causes of infertility and early pregnancy loss, which is linked to other pregnancy complications (8–12). Attainment of human endometrial receptivity occurs in the mid-secretory phase (MS) after sufficient time and concentration of progesterone exposure as seen in other placental mammals (13–18). In women without ovaries, sequential treatment with estrogen followed by estrogen plus progesterone, without any other ovarian hormones, is sufficient to achieve high rates of successful implantation of embryos derived from donor oocytes (16, 19), highlighting the importance of hormone actions in mediating implantation.

Abnormal progesterone signaling leads not only to fertility issues but also a spectrum of gynecological diseases (20–22), emphasizing the criticality of progesterone signaling in maintaining normal uterine biology and initiating pregnancy. The impact of progesterone is mediated through its nuclear receptor – Progesterone Receptor (PGR), where binding of progesterone induces its conformational change. This leads to altered affinity for target DNA response elements, thereby influencing the gene expression network at the transcriptional level (23). Although many PGR-regulated genes have been identified in both animal model systems and human studies as important mediators of implantation, including Indian Hedgehog (*IHH*) (24–26), Krüpple-like Factor 15 (*KLF15*) (27, 28), Heart and Neural Crest Derivatives-expressed 2 (*HAND2*) (29), Bone Morphogenesis Protein 2 (*BMP2*) (30, 31), Homeobox gene *HOXA10* (28, 32, 33), and CCAAT/Enhancer-binding Protein β (*CEBPB*) (34–36). Yet, implantation failure remains a great challenge in both natural pregnancies and assisted reproductive interventions.

Additionally, epithelial aspects of PGR actions are important, sometimes underappreciated determinants of implantation and pregnancy outcome. Endometrial epithelial cells line the uterine lumen and glands, with the latter derived from the former (37, 38). The endometrial epithelium undergoes dramatic cellular and molecular changes common to both mice and humans during the WOI, including adhesion mechanisms enabling the attachment of embryo to the luminal epithelium (39, 40), alterations in nuclear pore complex presentation (41), downregulation of the Serum and Glucocorticoid Regulated Kinase 1 (SGK1) (42), apoptotic cascade (43, 44), and expression of epithelial-specific receptivity markers (45). The glandular epithelium further facilitates implantation via the production of Leukemia Inhibitory Factor (LIF), a critical factor in embryo-uterine communication during WOI (46–48). Elaborate cross-talk exists between the endometrial epithelium and stroma that is indispensable for allowing implantation, adding further complexity to the regulatory mechanisms governing pregnancy establishment. Although animal model systems and *in vitro* cultured cells have proven instrumental in advancing our knowledge in reproductive functions, the high rate of implantation failure remain a challenge (49). The aim of this study is to use a single, comparative, human-derived, *ex vivo* analysis to examine the dynamics of PGR action during the WOI. We employed ChIP-seq technique to explore the modification of PGR binding landscape during the P to MS transition in human endometrial samples. Additionally, parallel RNA-sequencing analysis enabled the identification of differentially regulated genes, which allowed us to identify the subset of PGR-bound genes with altered mRNA abundance and hence relevance in regulating implantation and decidualization. Epithelial-specific RNA-sequencing allowed more precise assessment of the endometrial epithelial transcriptomic network, providing a deeper understanding of the dynamic transformation in the endometrium during the WOI.

## Results

### Characterization of PGR binding trend during the P and MS phases

To gain insights into the transcriptional regulatory function of PGR during the peri-implantation period, the physical association of PGR with DNA was assessed by PGR chromatin immunoprecipitation coupled to massively parallel DNA sequencing (ChIP-seq) using human endometrial biopsies from the P and MS phases. We identified over 10,000 genomic intervals (defined as a stretch of DNA sequence identified as exhibiting statistically significant PGR binding) as PGR bound in the endometrium. Analysis using the Peak Annotation and Visualization tool showed that majority of the PGR binding occurred within the intronic, intergenic, 5’ UTR and upstream region relative to the gene body, with no significant alteration in PGR binding preference to these categories between the two phases (Fig. 1. A).

**Figure 1.**
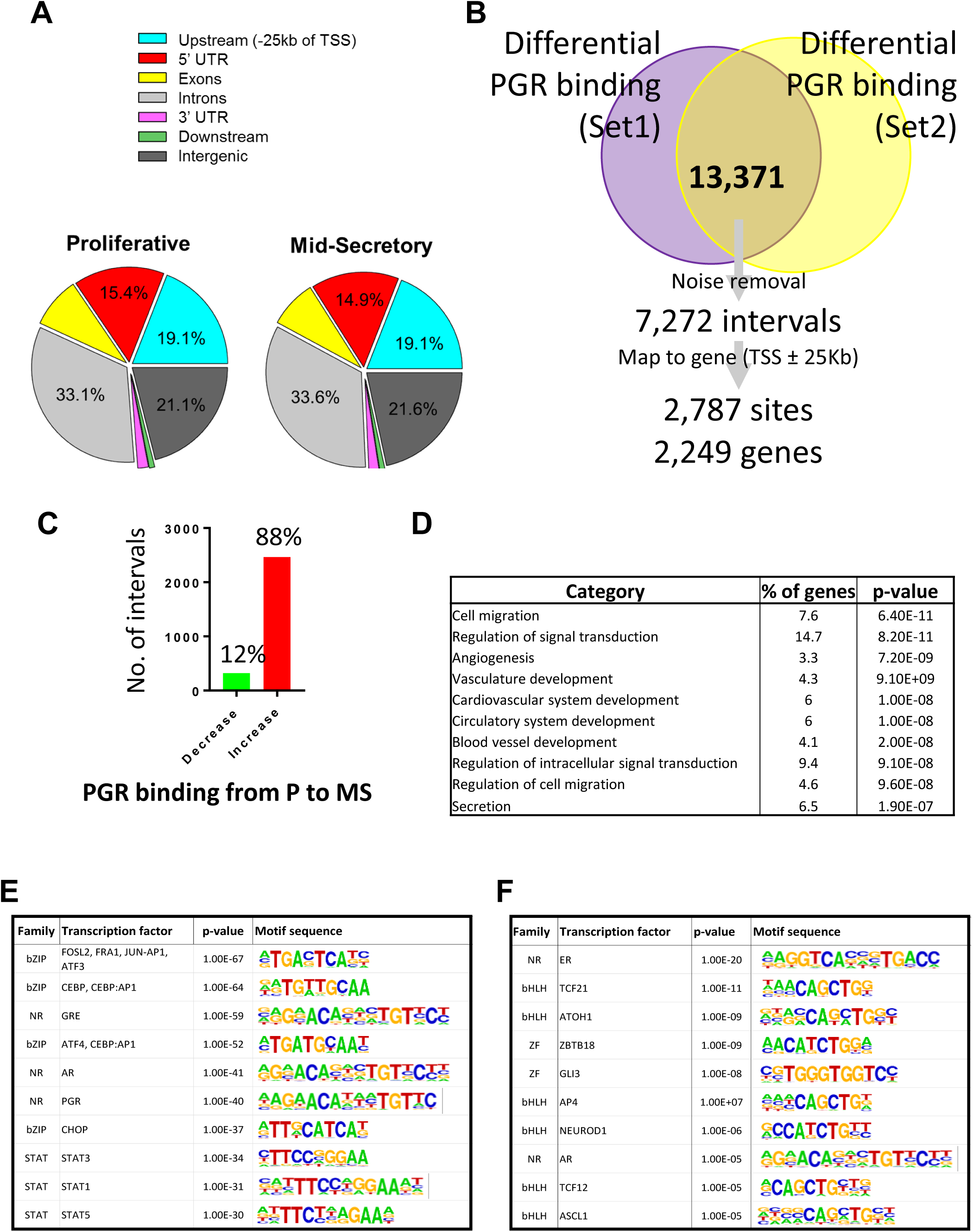
Genome wide PGR binding identified by ChIP-seq in endometrial tissue of fertile women during the proliferative and mid-secretory phases. (A). Distribution of PGR binding in the genome relative to the gene body during the P and MS phase, as analyzed by PAVIS. (B). Paired analysis was employed to identify differential PGR bound (DPRB) genomic intervals, where differential PGR binding was calculated for each of set1 and set2 (refer to Materials and Methods: *Chromatin immunoprecipitation sequencing (ChIP-seq) and qRT-PCR (ChIP-qPCR)*). The DPRB DNA common to both batches were defined as the real differential PGR bound sites. A total of 2,787 PGR bound regions were found to be in proximity of 2,249 genes (TSS ± 25 kb). (C). The percentage of total DPRB sites that showed increased (red) or decreased (green) PGR binding transitioning from P to MS. (D). Gene Ontology functional annotation showing enriched biological functions associated with DPRB genes (defined as DPRB within 25 kb of transcriptional start sites), as analyzed by the online bioinformatic tool DAVID. (E). Transcription factor binding sites enrichment in MS-gain intervals, as identified by the HOMER software. (F). Transcription factor binding sites enrichment in MS-loss intervals, as identified by the HOMER software.

Then, we characterized the PGR binding dynamics by identifying intervals with consistent or differential PGR binding (DPRB). Collectively, we analyzed two sets of samples each containing a P and MS pair. To circumvent the batch variation observed between the two sets of samples, we defined the consistent/constitutive PGR binding sites as those with PGR binding during both P and MS, where the read counts were not significantly different between P and MS in either one or both batches. For the DPRB intervals, we first analyzed each set independently to identify differential PGR binding sites, and only those DPRB common to both datasets were considered for additional analyses. In total, we identified 12,469 genomic sites with consistent PGR binding in proximity to 11,058 genes (Supplemental Table 1 (50)); and 2,787 genomic sites with altered PGR binding in proximity to 2,249 genes (Fig. 1. B, Supplemental Table 2 (50)). There were 2,466 intervals with increased PGR binding in proximity to 1,966 genes (88%) and 321 intervals with decreased PGR binding in proximity to 307 genes (12%, Fig. 1. C), and 423 genes were found with multiple differential PGR binding intervals in proximity.

Amongst the identified DPRB intervals, many were found in proximity to known PGR-regulated genes previously reported in both humans and mice, including FK506 Binding Protein 5 (*FKBP5*) (28), Indian Hedgehog (*IHH*), Insulin Receptor Substrate 2 (*IRS2*) (51), *CASP8* and FADD Like Apoptosis Regulator (*CFLAR*) (52), FOS Like 2 AP-1 Transcription Factor Subunit (*FOSL2*) (28), Perilipin 2 (*PLIN2*), Basic Leucine Zipper ATF-Like Transcription Factor (*BATF*) and Baculoviral IAP Repeat Containing 5 (*BIRC5*, Supplemental Table 2 (50)) (21). In addition, many known decidualizing and implantation mediators were found with constitutive PGR binding, including Forkhead Box Protein O1 (*FOXO1*) (53), Homeobox A10 (*HOXA10*) (53), Heart And Neural Crest Derivatives Expressed 2 (*HAND2*) (2), Cysteine Rich Angiogenic Inducer 61 (*CYR61*) (28) and Sex Determining Region Y-Box 17 (*SOX17*, Supplemental Table 1 (50)) (54, 55). The biological impact of PGR transcriptional activity during the P to MS phase was determined by examining the functional profile associated with the DPRB genes using the DAVID Bioinformatics Database (56, 57), and selected enriched pathways are shown in Table 1. Enrichment was observed in pathways regulating insulin resistance, focal adhesion, complement and coagulation cascades, cytokine-cytokine receptor interactions, ECM receptor interaction, apoptosis, as well as various signaling pathways including chemokines, Ras, FOXO, Prolactin, AMPK and Tumor Necrosis Factor (TNF). In addition, Gene Ontology functional annotation showed that the DPRB-associated genes are involved in the regulation of cell migration, signal transduction, angiogenesis, vasculature development and secretion (Fig. 1. D).

**TABLE 1.**
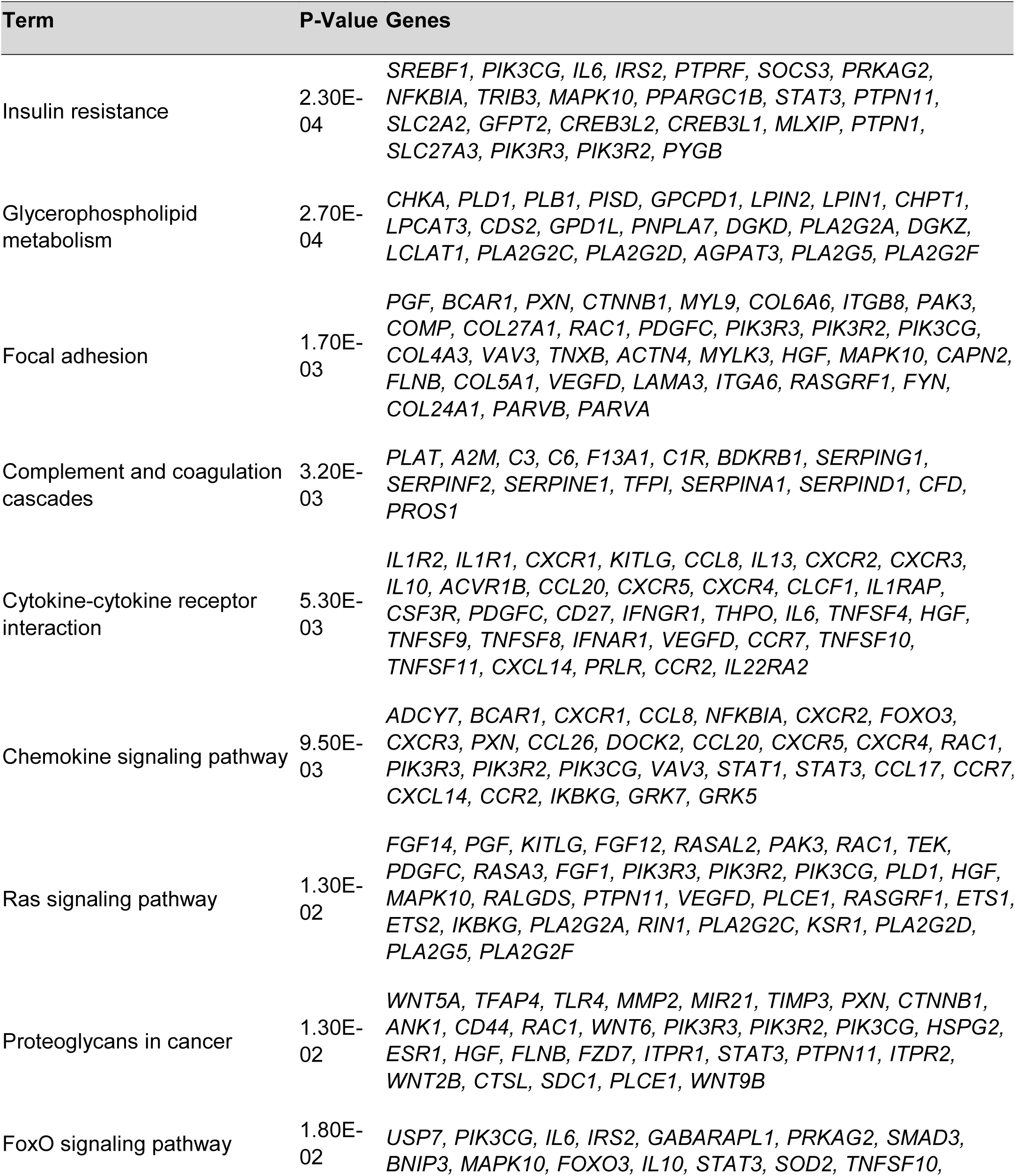
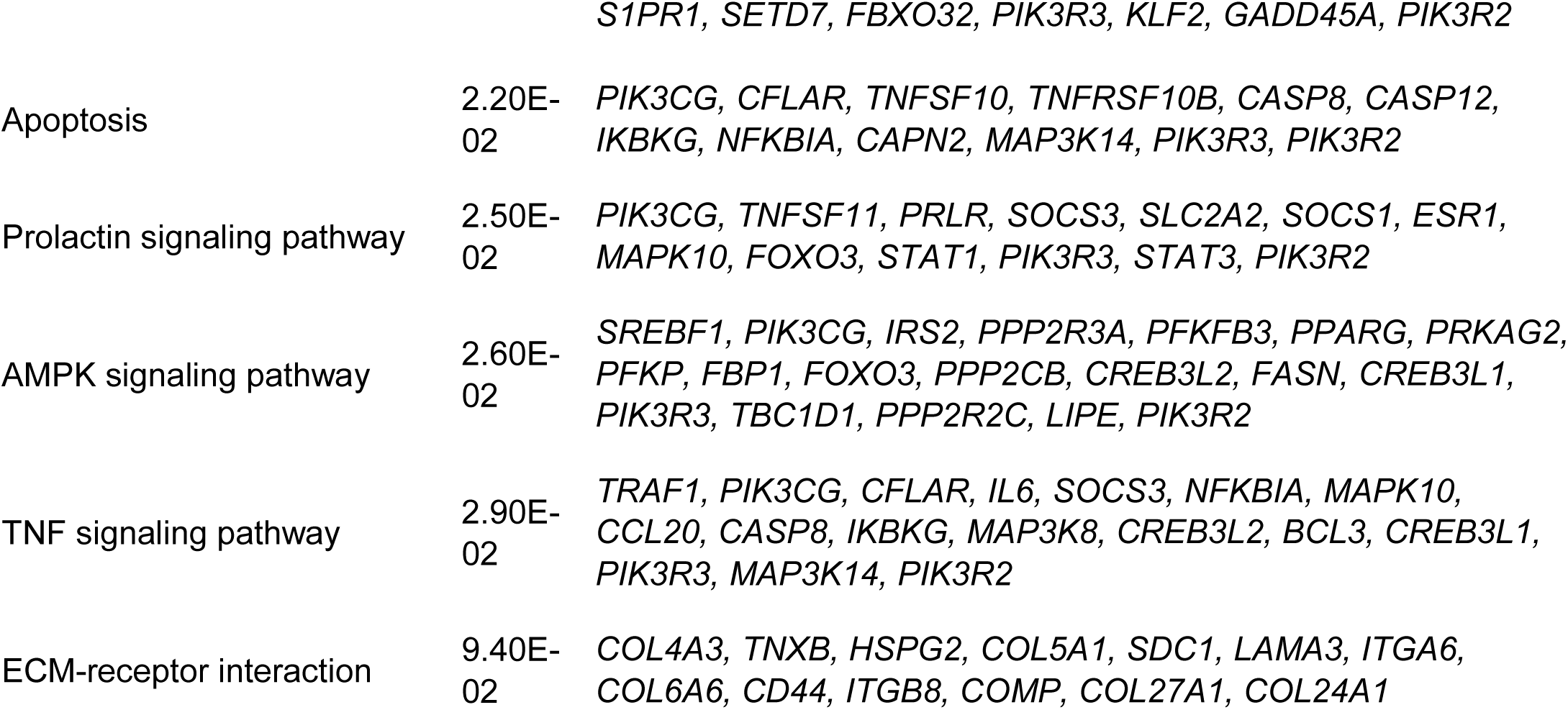
DAVID functional analysis using KEGG pathways for genes with differential PGR binding as determined by PGR ChIP-seq in the P and MS endometrium.

| Term | P-Value | Genes |
| --- | --- | --- |
| Insulin resistance | 2.30E-04 | *SREBF1, PIK3CG, IL6, IRS2, PTPRF, SOCS3, PRKAG2, NFKBIA, TRIB3, MAPK10, PPARGC1B, STAT3, PTPN11, SLC2A2, GFPT2, CREB3L2, CREB3L1, MLXIP, PTPN1, SLC27A3, PIK3R3, PIK3R2, PYGB* |
| Glycerophospholipid metabolism | 2.70E-04 | *CHKA, PLD1, PLB1, PISD, GPCPD1, LPIN2, LPIN1, CHPT1, LPCAT3, CDS2, GPD1L, PNPLA7, DGKD, PLA2G2A, DGKZ, LCLAT1, PLA2G2C, PLA2G2D, AGPAT3, PLA2G5, PLA2G2F* |
| Focal adhesion | 1.70E-03 | *PGF, BCAR1, PXN, CTNNB1, MYL9, COL6A6, ITGB8, PAK3, COMP, COL27A1, RAC1, PDGFC, PIK3R3, PIK3R2, PIK3CG, COL4A3, VAV3, TNXB, ACTN4, MYLK3, HGF, MAPK10, CAPN2, FLNB, COL5A1, VEGFD, LAMA3, ITGA6, RASGRF1, FYN, COL24A1, PARVB, PARVA* |
| Complement and coagulation cascades | 3.20E-03 | *PLAT, A2M, C3, C6, F13A1, C1R, BDKRB1, SERPING1, SERPINF2, SERPINE1, TFPI, SERPINA1, SERPIND1, CFD, PROS1* |
| Cytokine-cytokine receptor interaction | 5.30E-03 | *IL1R2, IL1R1, CXCR1, KITLG, CCL8, IL13, CXCR2, CXCR3, IL10, ACVR1B, CCL20, CXCR5, CXCR4, CLCF1, IL1RAP, CSF3R, PDGFC, CD27, IFNGR1, THPO, IL6, TNFSF4, HGF, TNFSF9, TNFSF8, IFNAR1, VEGFD, CCR7, TNFSF10, TNFSF11, CXCL14, PRLR, CCR2, IL22RA2* |
| Chemokine signaling pathway | 9.50E-03 | *ADCY7, BCAR1, CXCR1, CCL8, NFKBIA, CXCR2, FOXO3, CXCR3, PXN, CCL26, DOCK2, CCL20, CXCR5, CXCR4, RAC1, PIK3R3, PIK3R2, PIK3CG, VAV3, STAT1, STAT3, CCL17, CCR7, CXCL14, CCR2, IKBKG, GRK7, GRK5* |
| Ras signaling pathway | 1.30E-02 | *FGF14, PGF, KITLG, FGF12, RASAL2, PAK3, RAC1, TEK, PDGFC, RASA3, FGF1, PIK3R3, PIK3R2, PIK3CG, PLD1, HGF, MAPK10, RALGDS, PTPN11, VEGFD, PLCE1, RASGRF1, ETS1, ETS2, IKBKG, PLA2G2A, RIN1, PLA2G2C, KSR1, PLA2G2D, PLA2G5, PLA2G2F* |
| Proteoglycans in cancer | 1.30E-02 | *WNT5A, TFAP4, TLR4, MMP2, MIR21, TIMP3, PXN, CTNNB1, ANK1, CD44, RAC1, WNT6, PIK3R3, PIK3R2, PIK3CG, HSPG2, ESR1, HGF, FLNB, FZD7, ITPR1, STAT3, PTPN11, ITPR2, WNT2B, CTSL, SDC1, PLCE1, WNT9B* |
| FoxO signaling pathway | 1.80E-02 | *USP7, PIK3CG, IL6, IRS2, GABARAPL1, PRKAG2, SMAD3, BNIP3, MAPK10, FOXO3, IL10, STAT3, SOD2, TNFSF10,* |
|  |  | *S1PR1, SETD7, FBXO32, PIK3R3, KLF2, GADD45A, PIK3R2* |
| Apoptosis | 2.20E-02 | *PIK3CG, CFLAR, TNFSF10, TNFRSF10B, CASP8, CASP12, IKBKG, NFKBIA, CAPN2, MAP3K14, PIK3R3, PIK3R2* |
| Prolactin signaling pathway | 2.50E-02 | *PIK3CG, TNFSF11, PRLR, SOCS3, SLC2A2, SOCS1, ESR1, MAPK10, FOXO3, STAT1, PIK3R3, STAT3, PIK3R2* |
| AMPK signaling pathway | 2.60E-02 | *SREBF1, PIK3CG, IRS2, PPP2R3A, PFKFB3, PPARG, PRKAG2, PFKP, FBP1, FOXO3, PPP2CB, CREB3L2, FASN, CREB3L1, PIK3R3, TBC1D1, PPP2R2C, LIPE, PIK3R2* |
| TNF signaling pathway | 2.90E-02 | *TRAF1, PIK3CG, CFLAR, IL6, SOCS3, NFKBIA, MAPK10, CCL20, CASP8, IKBKG, MAP3K8, CREB3L2, BCL3, CREB3L1, PIK3R3, MAP3K14, PIK3R2* |
| ECM-receptor interaction | 9.40E-02 | *COL4A3, TNXB, HSPG2, COL5A1, SDC1, LAMA3, ITGA6, COL6A6, CD44, ITGB8, COMP, COL27A1, COL24A1* |

Despite the decrease in PGR expression during the MS phase (Supplemental Table 3 (50), see below), the global PGR binding trend was elevated as 88% of the intervals differentially bound by PGR exhibited increased binding during the MS phase (Fig. 1. C), which is likely due to the increased serum progesterone level in this phase of the cycle. To further explore enrichment of other transcription factor binding sites co-occupying the PGR binding intervals, the DPRB DNA motifs were analyzed by HOMER in two parts; those that showed elevated binding during MS (MS-gain) or reduced binding during MS (MS-loss). The MS-gain intervals, indeed, showed significant enrichment in PGR binding motif with a *p*-value of 1.00^-40^ (Fig. 1. E). MS-gain and MS-loss intervals exhibited distinct profiles of transcription factor binding site preferences, with *FOSL2*, *FRA1*, *JUN-AP1*, *ATF3* and *BATF* binding domains as top enriched sites in MS-gain intervals (Fig. 1. E). Nuclear Receptors *AR*, bZIP transcription factor *CHOP* and some STAT transcription factor members *STAT1*, *STAT3* and *STAT5* binding sites were also enriched in sites with increased PGR binding (Fig. 1. E). In contrast, enriched motifs in the MS-loss intervals included Estrogen Response Element (ERE), and binding domains for Transcription Factor 21 (*TCF21*), Atonal BHLH Transcription Factor 1 (*ATOH1*), Zinc Finger And BTB Domain Containing 18 (*ZBTB18*), as well as GLI Family Zinc Finger 3 (*GLI3*, Fig. 1. F). Of note, during the P to MS transition, PGR showed an increased preference for the Basic Leucine Zipper Domain (bZIP), as the MS-gain intervals belonged mainly to this class. On the other hand, preference for the Basic Helix Loop Helix (bHLH) and Zinc Finger (ZF) binding domains were lost during this phase transition, as the enriched motifs identified in the MS-loss intervals belonged mainly to these two classes. Thus, PGR’s effects on gene expression may be partially modulated through altered affinity for the different DNA responsive elements between the liganded and unliganded form.

### Transcriptional regulatory network of the P and MS endometrium

Whilst PGR has been widely studied in both humans and rodents and many direct and indirect target genes have been identified, a comprehensive analysis revealing its global regulatory function in the cycling human endometrium is still lacking. To fully characterize the functional relevance associated with PGR binding activities during the P to MS transition, we conducted RNA-seq on whole endometrium and incorporated the global gene expression profile into the ChIP-seq analyses during these two phases.

In total, we collected six P and five MS endometrial biopsies from which whole endometrial RNA was analyzed. This revealed a total of 14,985 expressed genes within the endometrium (FPKM > 1 in at least one of the two phases), whereby 14,303 and 14,156 were expressed in each of the P and MS phase, respectively. The transcriptomic profiles were subjected to hierarchical clustering and principal component analysis (PCA) as a measure of quality control. As shown in Supplemental Figure 1 (50). A, a distinct segregation was observed for the P- and MS-derived RNA expression profile, and this is further supported by the hierarchical clustering presented in the dendrogram shown in Supplemental Figure 1. B (50), where samples from the two stages clustered accordingly. This suggested that the samples were well-characterized according to stage and of appropriate quality.

Of the genes expressed in the endometrium, 4,576 were differentially expressed (DEGs, Supplemental Table 3 (50)) between the two phases (absolute fold change > 1.5; and adjusted *p* value < 0.05). In total, 2,392 genes showed increased expression while 2,184 were downregulated during MS. Several genes known to regulate uterine biology, decidualization and implantation were identified as DEGs including decidualizing markers IGF Binding Protein 1 (*IGFBP1*) and prolactin (*PRL*); hedgehog protein, Indian Hedgehog (*IHH*); transcription factors *FOXO1* and *GATA2*; Wnt signaling molecules *WNT4*, *WNT2*, *WNT5A* and their inhibitor *DKK1*; transcriptional repressor *ZEB1*; and extracellular matrix modulator *VCAN*. To interpret the biological impact of the DEGs during the P to MS transition, Gene Set Enrichment Analysis (GSEA) was performed to retrieve the functional profile associated with the DEGs (58). Consistent with current literature, elevated inflammatory response was identified as an enriched molecular function for the DEGs associated with the P to MS transition, as indicated by the positive enrichment in the TNFA-NFKB signaling axis, coagulation, allograft rejection, hypoxia, the complement cascade, interferon gamma response, IL6-JAK-STAT3 signaling and apoptosis (Table 2). On the other hand, the negatively enriched functions which represents repressed molecular pathways during MS showed significance in cell division regulatory mechanisms – including E2F targets, G2M checkpoint and mitotic spindle regulations (Table 2). The xenobiotic metabolism pathway was identified as one of the most positively enriched functions in the MS endometrium by both GSEA (Table 2, Fig. 2. A) and Ingenuity Pathway Analysis (data not shown). To validate the RNA-seq results, we examined expression of selected xenobiotic metabolism genes using RNA extracted from an independent set of endometrial biopsies (n = 6 for each of the P and MS phase), along with the expression of the decidualization markers *PRL* and *IGFBP1* to confirm the sample stages (Figs. 2. B and C). In accordance with the RNA-seq results (Fig. 2. D), the cytochrome P450 members *CYP2C18* and *CYP3A5*, solute carriers *SLC6A12* and *SLCO4A1*, and glucuronosyltransferase *UGT1A6* were all found to be upregulated during MS (Fig. 2. E). Further, glutathione S-transferase Mu genes (*GSTM1*, *GSTM3* and *GSTM5*), sulfotransferase *SULT1C4*, and solute carrier *SLCO2A1* were found to be repressed during the MS phase (Fig. 2. G) similarly to that observed with RNA-seq (Fig. 2. F).

**Figure 2.**
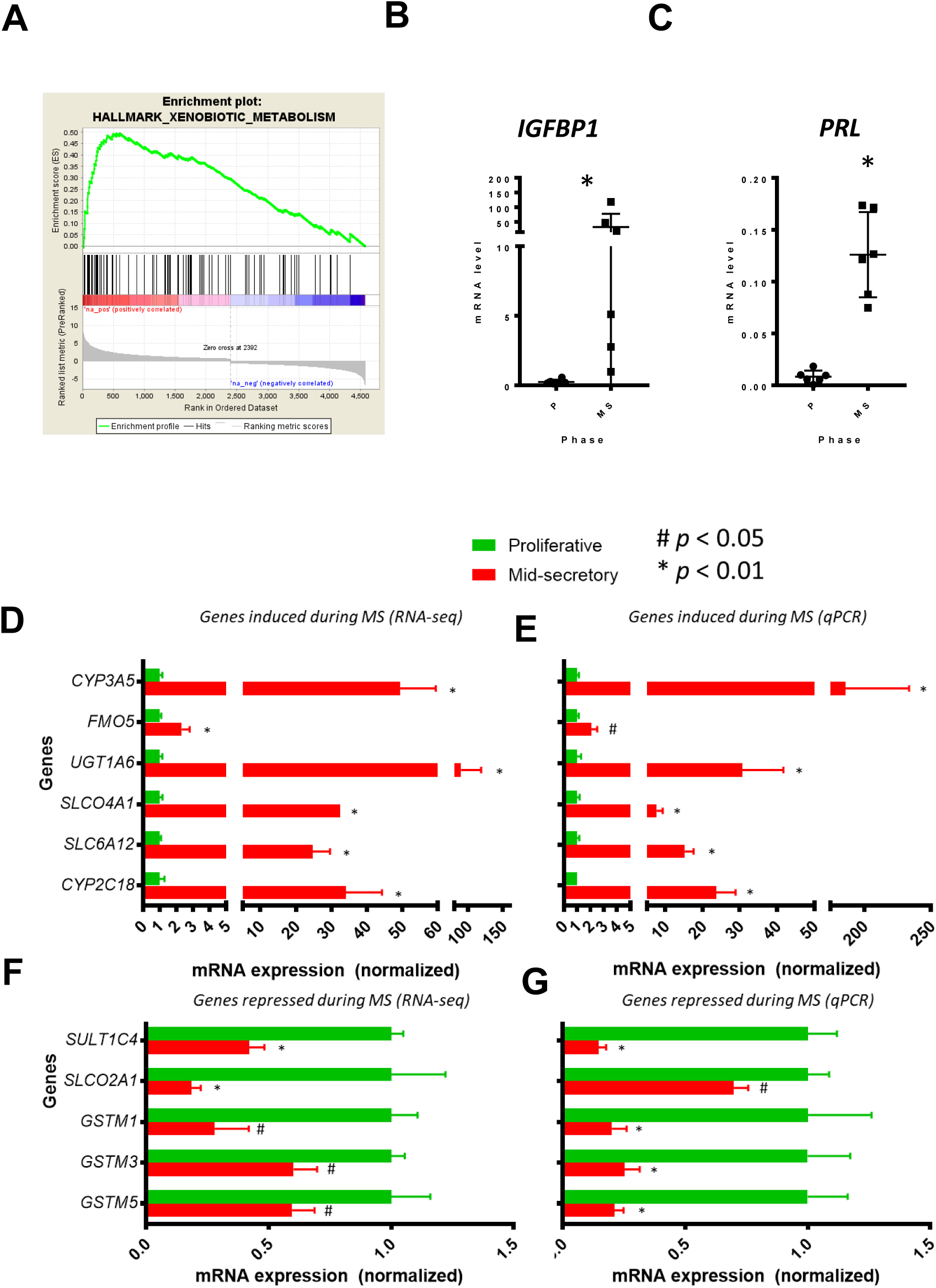
Endometrial gene expression profile during the proliferative and mid-secretory phases. (A). Gene Set Enrichment Analysis (GSEA) identified the xenobiotic metabolism pathway as significantly and positively enriched in the differentially expressed genes (DEGs), suggesting an increased activity in this pathway during MS. (B and C). Decidualization markers *IGFBP1* and *PRL* was examined by qRT-PCR using endometrial samples from independent patients to confirm stage of menstrual cycle. (D - G). Selected genes from the xenobiotic metabolism pathway were validated by qRT-PCR (E and G) using independent patient RNAs and presented in parallel with results from RNA-seq (D and F), n = 6, # *p* < 0.05 and * *p* < 0.01.

**TABLE 2.** Gene sets enrichment analysis of the 4,576 DEG in whole endometrium.

| Enriched gene sets | Enrichment | NES* | NOM p-val | FDR q-val |
| --- | --- | --- | --- | --- |
| TNFA SIGNALING VIA NFKB | Positive | 2.37 | 0 | 0 |
| XENOBIOTIC METABOLISM | Positive | 2.24 | 0 | 6.48E-04 |
| COAGULATION | Positive | 2.19 | 0 | 4.32E-04 |
| INFLAMMATORY RESPONSE | Positive | 2.15 | 0 | 3.24E-04 |
| COMPLEMENT | Positive | 1.83 | 0.00159236 | 0.012257015 |
| INTERFERON GAMMA RESPONSE | Positive | 1.77 | 0.0015456 | 0.017625704 |
| IL6 JAK STAT3 SIGNALING | Positive | 1.70 | 0.00980392 | 0.031572554 |
| APOPTOSIS | Positive | 1.54 | 0.02276423 | 0.08806576 |
| ANGIOGENESIS | Positive | 1.46 | 0.0970696 | 0.122301854 |
| E2F TARGETS | Negative | -3.12 | 0 | 0 |
| G2M CHECKPOINT | Negative | -3.04 | 0 | 0 |
| MITOTIC SPINDLE | Negative | -2.50 | 0 | 0 |
| MYC TARGETS V1 | Negative | -1.65 | 0.01590909 | 0.024345282 |
| DNA REPAIR | Negative | -0.96 | 0.4974359 | 0.69968206 |

\* NES = normalized enrichment score

### Functional profiling of DEGs with regulated PGR binding during P to MS

To search for the genes that are directly regulated by PGR and important in modulating implantation, we identified the genes that were both differentially expressed and differentially bound by PGR in the whole endometrium between the P and MS phases. Comparison of DEGs and DPRB gene lists revealed 653 genes common to both datasets (Fig. 3. A). The trend for PGR binding and altered gene expression during MS, as compared to P is summarized in Table 3 and graphically presented in Figure 3. B. This analysis found 87% of the genes showed increased PGR binding (572 out of 653), and 70% showed upregulation during the MS phase (454 out of 653). Interestingly, the majority of these genes showed a positive correlation between PGR binding change and transcriptional regulation, i.e. increased PGR binding was associated with increased gene expression and vice versa. Thus, PGR binding generally promotes rather than represses gene expression in the human endometrium (Fig. 3. C).

**Figure 3.**
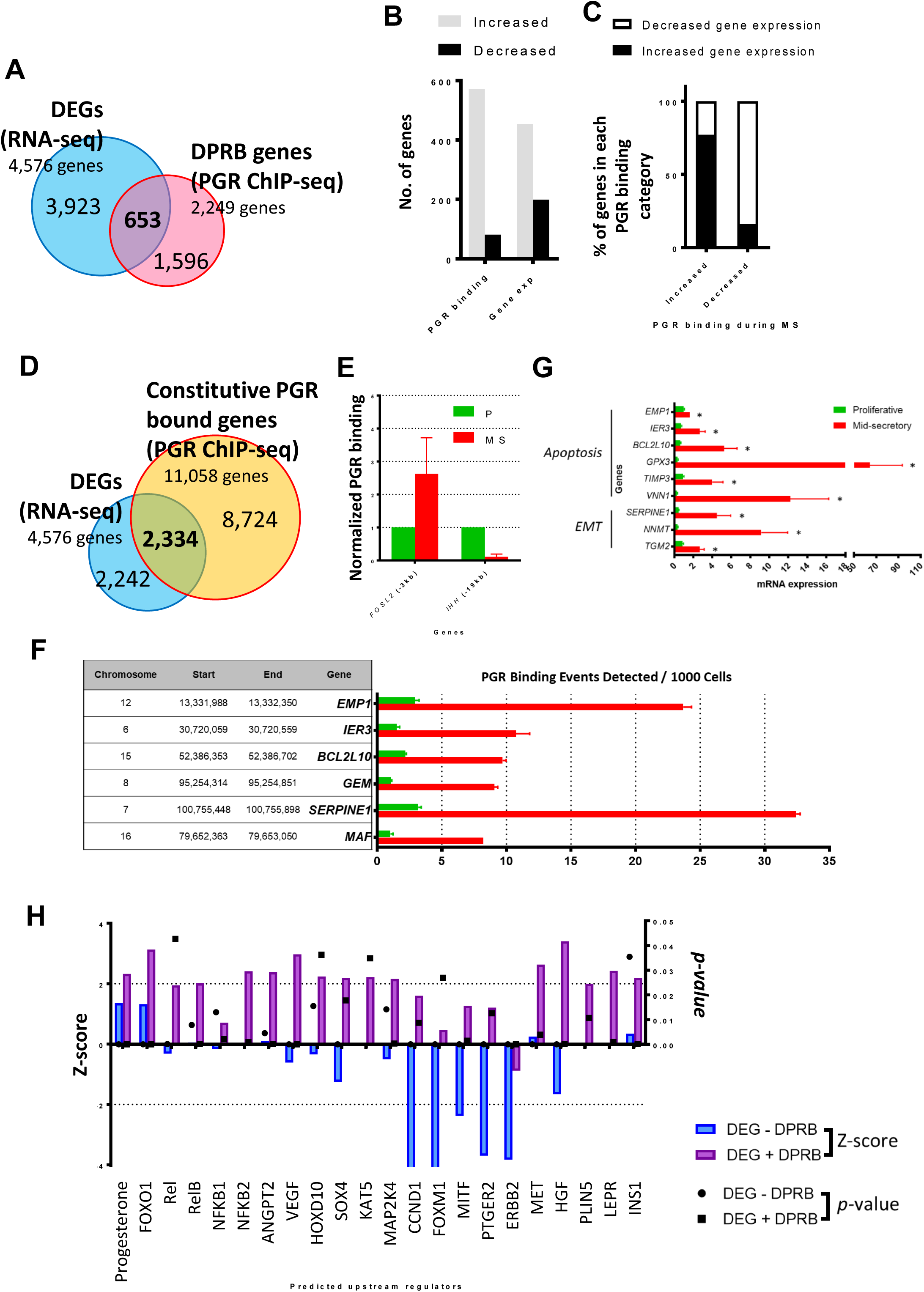
Identification of PGR regulated genes during the menstrual cycle. (A). Overlaying the genes with DPRB and differential expression identified 653 genes during the P to MS transition. (B). Number of genes showing increased and decreased PGR binding and expression in the endometrium during MS. (C). The percentage of genes showing increased or decreased expression with increased or decreased PGR binding from P to MS. (D). Overlaying the genes with proximal constitutive and PGR binding and differential expression identified 2,334 such genes during the P to MS transition. (E). PGR binding activity near two known target genes, *FOSL2* and *IHH* were examined by PGR ChIP-qPCR to confirm the phases of endometrial sample from which chromatin was obtained. qPCR was conducted in triplicates for each sample, and results shown are normalized to values from the P phase, n = 2 independent patients. (F). PGR occupancy was validated for selected genes from the xenobiotic metabolism, apoptosis and epithelial-mesenchymal transition (EMT) pathways using ChIP-qPCR. Experiments were performed using two sets of paired patient samples (each consisting of one P and one MS), and a representative result is shown. * *p* < 0.05. (G). Selected genes from the xenobiotic metabolism, apoptosis and EMT pathways were validated using qPCR, n = 6 and * *p* < 0.05. (H). Comparison of the upstream regulator activity (as indicated by the Z-score) for DEGs with and without differential PGR binding. Activity status (Z-score) is plotted on the left Y-axis (blue and purple bars, representing without DPRB and with DPRB, respectively), and significance (*p* value) is plotted on the right Y-axis (circle and square, representing without DPRB and with DPRB, respectively).

**TABLE 3.**
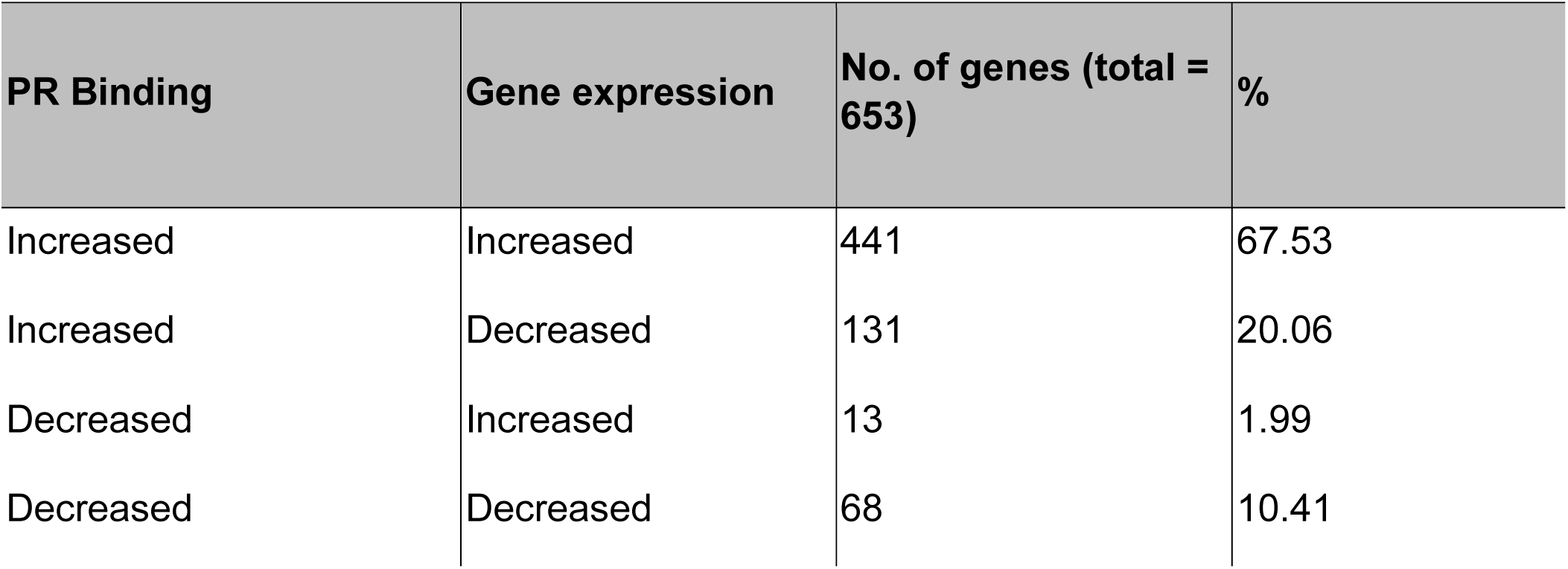
Genes with altered PGR binding and expression during P to MS transition.

| **PR Binding** | **Gene expression** | **No. of genes (total = 653)** | **%** |
| --- | --- | --- | --- |
| Increased | Increased | 441 | 67.53 |
| Increased | Decreased | 131 | 20.06 |
| Decreased | Increased | 13 | 1.99 |
| Decreased | Decreased | 68 | 10.41 |

The physiological function of PGR in regulating endometrial biology was next examined by elucidating the enriched functions associated with the PGR-regulated DEGs during the P to MS shift. The genes, along with fold change were submitted to GSEA to examine the enrichment of biological functions (Table 4). Enrichment was observed for a wide range of biological processes including inflammatory response signaling (coagulation, TNFA signaling via NFKB, complement, hypoxia, interferon gamma response), xenobiotic metabolism, epithelial mesenchymal transition (EMT), cell death regulation (apoptosis, p53 pathway), interleukin/STAT signaling, estrogen response, and MTORC1 response. Many of these biological functions were similarly identified using the DAVID Bioinformatic Database such as the regulation of cell death, inflammatory response, cytokine production, response to hormone and response to oxygen levels (Supplemental Table 4 (50)). Additionally, “secretion by cell” was identified as a regulated pathway by DAVID (*p* = 6.60E-5), supporting the validity of the secretory-phase derived gene expression profile. Other pathways identified by DAVID included cell migration, signal transduction, angiogenesis, leucocyte migration, nitric oxide biosynthetic processes, ECM disassembly, and various activities associated with lipid regulation and insulin response (Supplemental Table 4 (50)).

**TABLE 4.** Gene sets enrichment analysis of the 653 genes with differential expression and differential PGR binding.

| Enriched gene sets | Enrichment | NES | NOM p-val | FDR q-val |
| --- | --- | --- | --- | --- |
| COAGULATION | Positive | 1.85 | 0.001364257 | 0.0448207 |
| INFLAMMATORY RESPONSE | Positive | 1.59 | 0.037333332 | 0.2136788 |
| TNFA SIGNALING VIA NFKB | Positive | 1.59 | 0.026041666 | 0.1491825 |
| XENOBIOTIC METABOLISM | Positive | 1.57 | 0.03547963 | 0.1247853 |
| EPITHELIAL MESENCHYMAL TRANSITION | Positive | 1.55 | 0.038208168 | 0.1259948 |
| COMPLEMENT | Positive | 1.38 | 0.11479945 | 0.2892615 |
| APOPTOSIS | Positive | 1.37 | 0.092369474 | 0.2594335 |
| HYPOXIA | Positive | 1.31 | 0.1342711 | 0.3065951 |
| INTERFERON GAMMA RESPONSE | Positive | 1.29 | 0.17036012 | 0.302116 |
| ESTROGEN RESPONSE LATE | Positive | 1.15 | 0.31600547 | 0.4711447 |
| IL2 STAT5 SIGNALING | Positive | 1.13 | 0.31117022 | 0.4615895 |
| P53 PATHWAY | Positive | 1.12 | 0.32647464 | 0.4400889 |
| MTORC1 SIGNALING | Positive | 0.89 | 0.6025825 | 0.7500677 |
| ESTROGEN RESPONSE EARLY | Positive | 0.86 | 0.6555407 | 0.7511974 |
| IL6 JAK STAT3 SIGNALING | Positive | 0.76 | 0.7735584 | 0.8316847 |

Additionally, some genes known to regulate decidualization and implantation showed constitutive PGR binding during both phases (*FOXO1*, *HOXA10*, *HAND2*, *SOX17* and *CYR61*), suggesting that constitutive PGR binding may regulate endometrial functions. We thus examined the biological significance of the DEGs with constitutive PGR binding. Overlaying the constitutive PGR bound genes (Supplemental Table 1 (50)) and DEGs (Supplemental Table 3 (50)) identified 2,334 common genes (Fig. 3. D). The consistent PGR binding to these genes suggest that their altered expression is not regulated directly by altered PGR binding and may require input from other regulatory factors. Evaluation of the biological processes controlled by this group of genes showed primarily proliferative functions (cell cycle, cell division, nuclear division, DNA replication, Supplemental Table 5 (50)), and further analysis using GSEA confirmed that the proliferative function is repressed (Table 5, negatively enriched pathways). Additionally, TNFA signaling via NFKB, inflammatory response and hypoxia were identified as top positively enriched signaling pathways associated with this group of genes. Comparison of the functional profile defined by the DEGs that were differentially (Table 4) or constitutively (Table 5) bound by PGR showed some common signaling pathways involving both groups of genes. However, DEGs with DPBR appear to engage more specifically with functions including coagulation, EMT, estrogen response and apoptosis.

**Table 5.** Gene sets enrichment analysis of the 2,334 DEGs in whole endometrium with constitutive PGR binding.

| Enriched gene sets | Enrichment | NES | NOM p-val | FDR q-val |
| --- | --- | --- | --- | --- |
| TNFA SIGNALING VIA NFKB | Positive | 2.27366 | 0 | 0.002744444 |
| INFLAMMATORY RESPONSE | Positive | 2.2703116 | 0 | 0.001372222 |
| HYPOXIA | Positive | 2.1009197 | 0 | 0.004157759 |
| INTERFERON GAMMA RESPONSE | Positive | 1.9754444 | 0 | 0.01090216 |
| IL6 JAK STAT3 SIGNALING | Positive | 1.8338909 | 0.017621145 | 0.02559644 |
| XENOBIOTIC METABOLISM | Positive | 1.6081412 | 0.025882352 | 0.10718091 |
| KRAS SIGNALING UP | Positive | 1.5911685 | 0.03671706 | 0.10251326 |
| UV RESPONSE DN | Positive | 1.5904794 | 0.015521064 | 0.09035362 |
| COMPLEMENT | Positive | 1.5408636 | 0.04405286 | 0.10960495 |
| E2F TARGETS | Negative | -3.255928 | 0 | 0 |
| G2M CHECKPOINT | Negative | -3.1368256 | 0 | 0 |
| MITOTIC SPINDLE | Negative | -2.8043678 | 0 | 0 |
| HEDGEHOG SIGNALING | Negative | -1.9978325 | 0 | 0.002890576 |
| EPITHELIAL MESENCHYMAL TRANSITION | Negative | -1.4042959 | 0.06571936 | 0.14482453 |

To authenticate the ChIP-seq results and the regulatory role of PGR, PGR-chromatin association was evaluated for selected genes from the apoptosis and EMT pathways, both of which are known to regulate receptivity. In addition, we examined PGR binding near the MAF bZIP Transcription Factor (*MAF*), a regulator of the xenobiotic metabolism pathway shown earlier to be positively enriched during MS. Two known PGR-regulated genes in the human endometrial cells, *IHH* and *FOSL2* were first validated and confirmed to show increased (*FOSL2*) and decreased (*IHH*) PGR binding during the MS phase (Figs. 3. E). Apoptosis regulating genes Epithelial Membrane Protein 1 (*EMP1*), Immediate Early Response 3 (*IER3*), and B-Cell CLL/Lymphoma 2 Like 10 (*BCL2L10*), as well as EMT mediators GTP Binding Protein Overexpressed In Skeletal Muscles (*GEM*) and Serpin Family E Member 1 (*SERPINE1*), all displayed elevated PGR binding during the MS phase indicated by independent ChIP-qPCR analysis (Fig. 3. F). Additionally, independent qPCR analysis revealed the elevated transcription of apoptotic modulators (*EMP1*, *IER3* and *BCL2L10*) and the EMT regulator *SERPINE1*. Other genes regulating these two pathways were also found to be transcriptionally regulated, including Glutathione Peroxidase 3 (*GPX3*), Tissue Inhibitor Of Metalloproteinases 3 (*TIMP3*), Vanin 1 (*VNN1*), Nicotinamide N-Methyltransferase (*NNMT*) and Transglutaminase 2 (*TGM2*, Fig. 3. G).

To identify potential regulators associated with PGR, we next used IPA to predict for activity of upstream regulators based on the 653 common genes (DEG + DPRB), and DEGs without differential PR binding (DEG – DPRB, 3,923 genes), and upstream regulators were compared. This comparison showed a higher Z-score for both progesterone and FOXO1 (a known co-factor of PGR) in the regulation of the DEG + DPRB genes compared to the DEG – DPRB genes (Fig. 3. H), confirming that this group of genes is more closely associated with the progesterone-PGR signaling. Amongst the upstream regulators predicted for each gene set, the inflammation associated transcription factor NFKB family including REL, RELB and NFKB2 all possessed a stronger activation score in the DEGs + DPRB (Fig. 3. H), suggesting enhanced activity based on the altered gene expression network. In addition to NFKB, the angiogenic modulators ANGPT2 and VEGF, developmental regulators HOXD10 and SOX4, histone modifier KAT5 and the kinase MAP2K4 were all regulators predicted to have a higher activation score in regulating the group of genes with differential PGR binding. Interestingly, the cell cycle regulator CCND1, transcriptional regulators FOXM1 and MITF, prostaglandin receptor PTGER2 and the kinase protein ERBB2 were all predicted to be strongly inhibited in the regulation of DEG - DPRB, but Z-score prediction suggest that those factors were not inhibited in the regulation of the DEGs + DPRB. This suggests that although PGR may not directly inhibit these factors, they may engage with PGR in a co-operative manner to regulate the downstream gene expression network. Moreover, the MET-HGF receptor ligand pair as well as fat metabolism modulators PLIN5, LEPR and Insulin I were all found with increased activity in regulating the DEGs + DPRB, suggesting that these signaling axes are also associated with PGR function in the cycling human uterus. Interestingly, although Insulin (*INS*) itself was not transcriptionally regulated during the P to MS cycle, its cognate receptor Insulin Receptor (*INSR*) showed strong transcriptional induction (Supplemental Table 3 (50)). Additionally, many genes known to be regulated by insulin including *TIMP3* (Fig. 3. G, Supplemental Table 3 (50)), *SOD2*, *SOCS3*, *PRLR* and *MMP2* all showed elevated mRNA expression in the MS endometrium (Supplemental Table 3 (50)).

### Epithelial transcriptome in the cycling endometrium

To further understand the complexity of the cycling uterus, we assessed transcriptional changes in the epithelial lining of the endometrium. As the endometrium consists of a complex and dynamically changing set of cells, gene expression profiles derived from whole endometrial biopsies often overlook alterations of specific cell types. Four P and five MS endometrial samples were obtained, from which the luminal and glandular epithelial RNA were extracted and subjected to RNA-seq analysis. Principal component analysis (PCA) and hierarchical clustering found good segregation of the gene expression profile derived from two differently staged samples (Supplemental Fig. 2. A and B (50)). In the epithelium, we found a comparable number of genes expressed to that of the whole endometrium, with 14,502 genes and 13,993 genes transcriptionally active during the P and MS phase, respectively. The same threshold for identifying DEGs in the whole endometrium was applied to the epithelium-expressed genes, with which 3,052 epithelial-specific DEGs were found (epi-DEGs, Supplemental Table 6 (50)). Of those, 57% (1,764) showed elevated transcription and 43% (1,288) was transcriptionally repressed during the MS phase. Functional enrichment analysis of the epi-DEGs using GSEA showed a positive enrichment for the genes encoding components of the apical junction complex (Table 6), a molecular process important in defining the polarity of the epithelium and hence supports the authenticity of the gene expression profile obtained from an epithelial origin. Most of the pathways identified for the epithelium, whether positively or negatively enriched, were principally similar to that of the whole endometrium, with enrichment in pathways regulating immune responses including coagulation, complement, TNFA signaling via NFKB, apoptosis, as well as xenobiotic metabolism. On the other hand, cell division related processes including E2F regulated cell cycle, G2M checkpoint, mitotic spindle and DNA repair were negatively enriched (Table 6).

**TABLE 6.** Gene sets enrichment analysis of the 3,052 DEGs in the epithelium.

| Enriched gene sets | Enrichment | NES | NOM p-val | FDR q-val |
| --- | --- | --- | --- | --- |
| COAGULATION | Positive | 2.4593391 | 0 | 0 |
| COMPLEMENT | Positive | 2.3820252 | 0 | 0 |
| TNFA SIGNALING VIA NFKB | Positive | 2.260513 | 0 | 0 |
| INFLAMMATORY RESPONSE | Positive | 2.2356772 | 0 | 0 |
| XENOBIOTIC METABOLISM | Positive | 2.1790328 | 0 | 0.000179012 |
| HYPOXIA | Positive | 1.9205347 | 0 | 0.008392723 |
| APOPTOSIS | Positive | 1.8877827 | 0.001414427 | 0.009475978 |
| INTERFERON GAMMA RESPONSE | Positive | 1.8587548 | 0 | 0.011746863 |
| KRAS SIGNALING UP | Positive | 1.8500544 | 0.004172462 | 0.011368177 |
| EPITHELIAL MESENCHYMAL TRANSITION | Positive | 1.6994203 | 0.004178273 | 0.03300058 |
| ANGIOGENESIS | Positive | 1.4216607 | 0.09548611 | 0.15824698 |
| APICAL JUNCTION | Positive | 1.2970207 | 0.15987934 | 0.27779698 |
| P53 PATHWAY | Positive | 1.2064549 | 0.21958457 | 0.38236877 |
| E2F TARGETS | Negative | -<br>3.7750194 | 0 | 0 |
| G2M CHECKPOINT | Negative | -<br>3.4905815 | 0 | 0 |
| MYC TARGETS V1 | Negative | -<br>2.5305622 | 0 | 0 |
| MITOTIC SPINDLE | Negative | -<br>2.4694543 | 0 | 0 |
| ESTROGEN RESPONSE LATE | Negative | -<br>1.8183266 | 0 | 0.009814334 |
| DNA REPAIR | Negative | -<br>1.0963912 | 0.32970026 | 0.43156412 |
| MTORC1 SIGNALING | Negative | -<br>0.7685945 | 0.8393939 | 0.88358915 |
| ESTROGEN RESPONSE EARLY | Negative | -<br>0.7567437 | 0.8778878 | 0.8235289 |

### Functions specific to the endometrial epithelium during implantation

To tease out the epithelial specific molecular events, we next compared the whole endometrium DEG with the epi-DEG to identify DEGs that are unique to the epithelium. In total, 2,411 common genes were found, representing those that show differential expression in both the whole endometrium and epithelium (Fig 4. A, Supplemental Tables 3 and 6 (50)). Of those, 2,394 genes showed the same transcriptional change between the two compartments, and 17 genes, although identified as “common” DEGs, exhibited the reversed change in mRNA level between the two compartments. Altogether with the 641 genes which were exclusively regulated in the epithelium, a total of 658 genes which were “specifically” regulated in the epithelium was found. Canonical pathways regulated by this group of genes were assessed using IPA and ranked according to significance in Table 7. Synthesis of glycosaminoglycans, including dermatan sulfate and chondroitin sulfate, as well as cholesterol biosynthesis were the most significant pathways identified (-Log *p* value > 3). Osteoarthritis pathway, cholecystokinin/gastrin mediated signaling, IL8 signaling and TGFB signaling were all significant pathways with a positive Z-score, suggesting increased activity during MS in the endometrial epithelium. On the other hand, PTEN signaling was identified as significantly repressed in the MS epithelium (Table 7).

**Figure 4.**
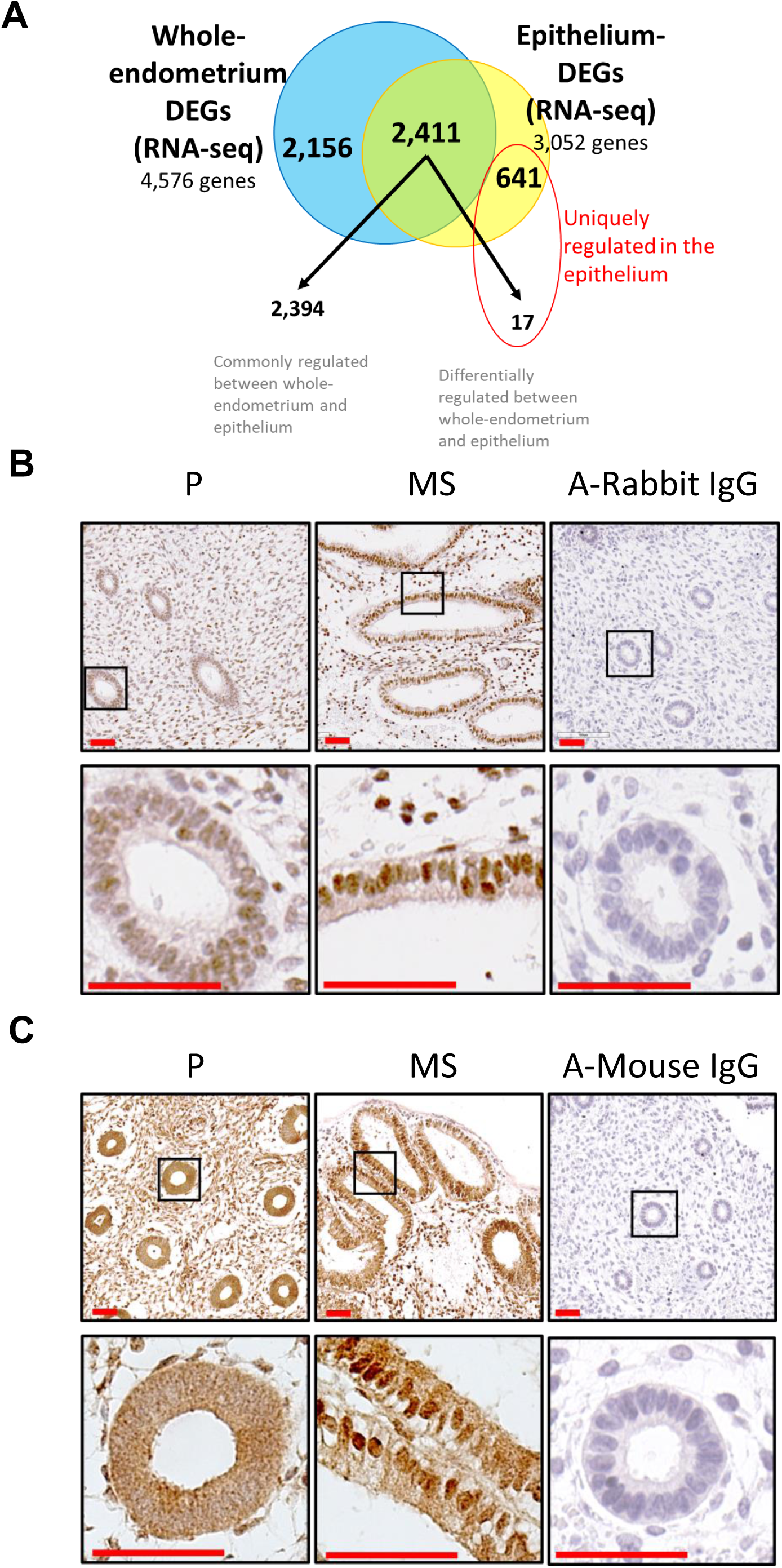
Epithelial functions during implantation and protein regulation of epithelial regulators IRF8 and MEF2C. (A). Comparison of DEGs derived from the epithelium to DEGs derived from the whole endometrium, with a total of 658 genes that were uniquely regulated in the epithelium. (B – C) Immunohistochemistry staining for IRF8 (B) and MEF2C (C) in human endometrial samples during P and MS. Results show that both proteins were expressed in both the epithelium, with increased levels of IRF8 and increased cytoplasmic-nuclear translocation of MEF2C during the MS phase. Experiment was conducted on three independent patients’ samples and a representative is shown, alongside the negative control stained with secondary antibody only.

**TABLE 7.**
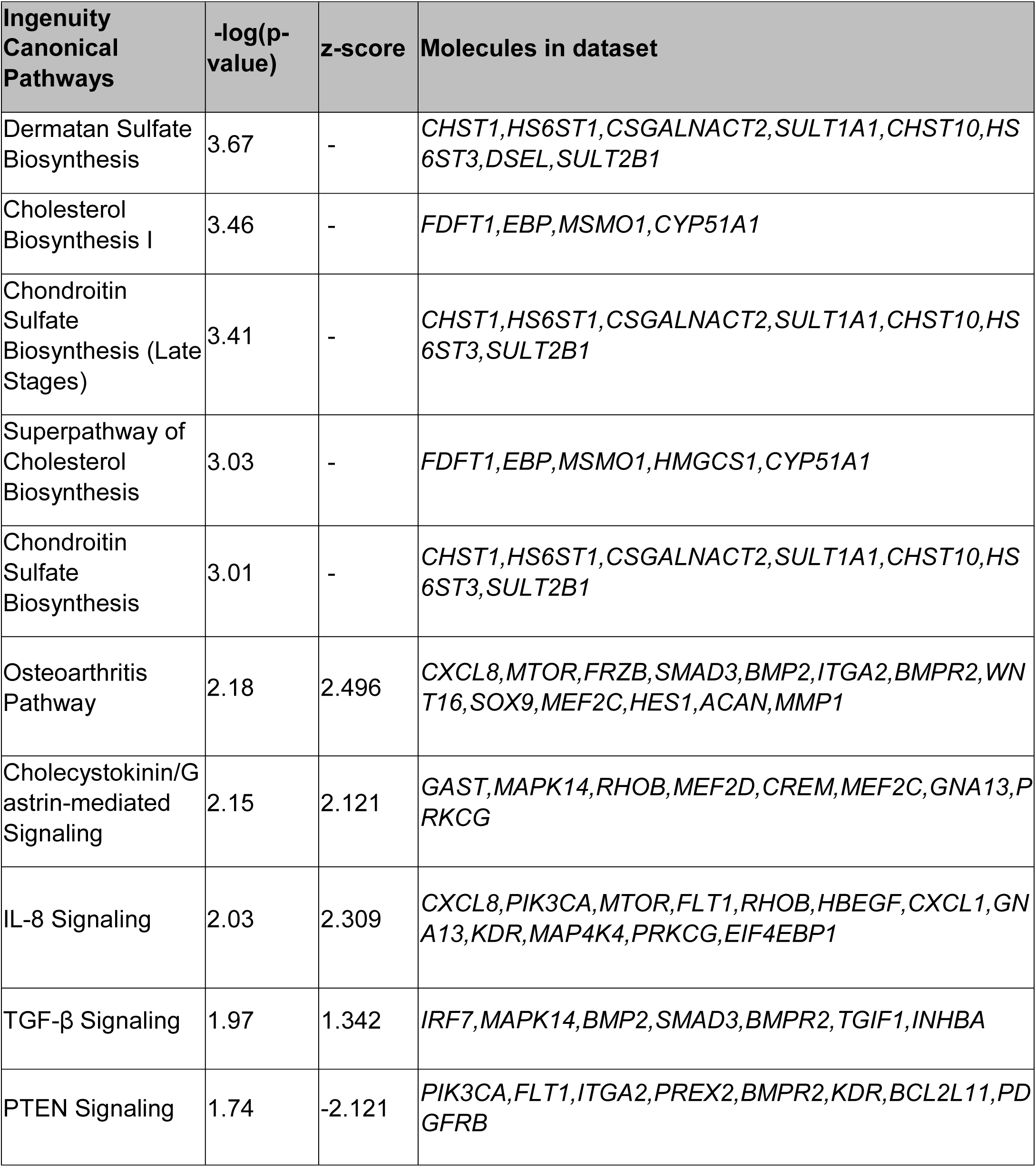
Canonical pathway analysis of the epithelium specific DEGs using IPA.

IPA was next used to predict for upstream regulator activities in the epithelium (Supplemental Table 7 (50)). As expected, both progesterone and PGR were identified as activated upstream regulators, with Z-score values of 2.269 and 3.812, and *p* values of 1.77E-46 and 1.29E-29, for progesterone and PGR, respectively. Estrogen Receptor Alpha (ESR1) was shown to be repressed while ESR2 was activated. Interestingly, RNA-seq results illustrated decreased expression of *ESR1* and upregulation of *ESR2* in the MS epithelium (Supplemental Table 6 (50)). The top activated regulators were cytokines including IL1B, TNF, IFNG, OSM and IL1A; as well as transcriptional regulators such as NUPR1, NFKB, SMARCA4 and CEBPA (Z-score > 5). Repressed regulators included transcription factors TBX2, TAL1; small GTPase RABL6, as well as the E1A Binding Protein P400 (EP400).

Lastly, to identify regulators with specific activities in the epithelium during the MS phase, we cross-compared the upstream regulators identified for the whole endometrium DEGs and epithelial DEGs. To ensure that the upstream regulators identified were meaningful and relevant, we compared only the regulators with *p-*values less than 0.05, and the numerical activation Z-score values greater than 1.5. This comparison yielded several regulator proteins with specific actions in the epithelium, of which selected are shown in Table 8. Amongst those were transcriptional regulators POU5F1, IRF5, IRF8 and FOXJ1; Myocyte Enhancer Factors family MEF2C and MEF2D; transmembrane receptors TLR5, IL1R1 and FCGR2A; kinase proteins MET and AURKB; the growth factor HBEGF; the CYP27B1 enzyme and Wnt ligand WNT7A; as well as the Notch ligand DLL4. Interestingly, *MEF2C*, *MEF2D*, *IRF8*, *FOXJ1*, *HBEGF*, *CYP27B1* and *DLL4* were found to be uniquely regulated in the epithelium, where either transcriptional regulation was not detected in the whole endometrium or showed a different pattern of gene expression during the P to MS transition. In addition, *HBEGF* was detected at very low levels as indicated by an average FPKM value of 2.63 in the whole endometrium; compared to 19.26 in the epithelium (data not shown), suggesting that the transcription of this gene is enriched in the epithelial cells during the WOI. We examined the protein expression of two epithelial-specific regulators IRF8 and MEF2C using formalin-fixed and paraffin-embedded endometrial biopsies from independent patients. As shown in Figure 4. B, IRF8 is expressed in both stromal and epithelial cells and exhibited elevated protein expression during the MS phase. MEF2C, on the other hand, did not show substantial increase in staining intensity, but displayed a robust cytoplasmic-to-nuclear translocation from the P to MS stage in the glandular epithelium (Fig. 4. C). These results suggest that both IRF8 and MEF2C, two proteins previously unreported to have a role in implantation are regulated both at the mRNA and protein level during the peri-implantation phase of the menstrual cycle in the epithelium, and hence may have important functions in the implantation-phase endometrium.

**TABLE 8.** Upstream regulators with specific actions in the epithelium (identified using IPA).

| Upstream Regulator | Log2(FC) in epithelium | Molecule Type | Activation z-score | p-value of overlap |
| --- | --- | --- | --- | --- |
| POU5F1 | 1.696 | transcription regulator | 2.429 | 3.93E-06 |
| IRF5 | 1.526 | transcription regulator | 2.266 | 2.04E-04 |
| MEF2C | 1.098 | transcription regulator | 1.835 | 6.15E-03 |
| MEF2D | 1.014 | transcription regulator | 2.478 | 1.02E-03 |
| IRF8 | 1.199 | transcription regulator | 1.608 | 3.35E-05 |
| FOXJ1 | -1.613 | transcription regulator | -1.96 | 3.52E-02 |
| TLR5 | 1.34 | transmembrane receptor | 3.062 | 7.08E-04 |
| IL1R1 | 1.745 | transmembrane receptor | 2.872 | 2.27E-02 |
| FCGR2A | 2.025 | transmembrane receptor | 1.544 | 1.30E-04 |
| MET | 1.993 | kinase | 2.054 | 1.26E-09 |
| AURKB | -3.136 | kinase | -2.132 | 2.99E-03 |
| HBEGF | 1.739 | growth factor | 1.754 | 1.94E-05 |
| WNT7A | -2.761 | Wnt ligand | -1.98 | 2.60E-02 |
| CYP27B1 | -1.773 | enzyme | -1.51 | 9.40E-03 |
| DLL4 | 1.067 | other | 2.119 | 1.74E-05 |

## Discussion

Here, we investigated the cycling human endometrium at the molecular level with two major aims in mind. First, to gain better understanding of the human endometrial signaling pathways and molecular events controlled by PGR during the P to MS transition. Combining the PGR cistromic and whole endometrium transcriptomic profile allowed the identification of genes with both proximal PGR binding and transcriptional regulation during the WOI. Second, we examined the gene expression profile using RNA derived from the whole endometrium or from the epithelium, including both luminal and glandular. Comparison of the two expression profiles delineated a more sophisticated and compartment specific transcriptional network. The latter has remained a challenging task and for this reason, the endometrium has often been examined as a whole when conducting *in vivo* studies.

### The biological significance of PGR transcriptional activity during the WOI

Using PGR ChIP-seq, we obtained a genome wide DNA-binding blueprint of PGR in the endometrium at the P and MS phases. Comparison of the two identified DEGs with constitutive or regulated PGR binding in proximity during this period. Using the motif finding tool HOMER, we found a distinguishing difference in PGR binding preference from P to MS. While sites with increased PGR bindings at MS were predominantly co-occupied by bZIP and STAT transcription factors, sites with reduced PGR binding during MS were shared by bHLH and ZF transcription factors. This finding may suggest a mechanism of regulation for PGR transcriptional activity whereby its preference for certain DNA motifs is gained or lost during different phases of the menstrual cycle. Alternatively, the association of PGR to these DNA motifs may not be a direct one, but rather through interaction with other transcription factors which then associate with the promoter region. The changes in DNA motifs detected based on altered PGR binding could in turn suggest a change in PGR preference for different transcription factors rather than different DNA motifs. Indeed, PGR is known to control gene expression in this way through transcription factors such as SP1 and AP1 in human endometrial cells and mammary cells (28, 59, 60).

A more comprehensive landscape of PGR biological impact was achieved by comparing the whole endometrium derived DEGs to genes with DPRB in proximity to identify genes whose transcription is likely directly regulated during the menstrual cycle by PGR. We found 653 such genes, and analysis by GSEA identified many enriched pathways one of which is the metabolism of xenobiotics. To the best of our knowledge, our study is the first to identify xenobiotic metabolism as a PGR regulated pathway in the cycling endometrium. Xenobiotics are conventionally defined as entities foreign to a cell or tissue such as drugs and pollutants, although it can also refer to entities found at levels greater than considered norm. Xenobiotic metabolism hence refers to the modification of these entities which in turn allows their systemic removal. Genes involved in this pathway are broadly categorized into 3 phases: phase 1 and phase 2 enzymes increase the solubility of the xenobiotics by introducing polar moieties and conjugating to endogenous hydrophilic molecules; and phase 3 genes encode transporters which then traffic the xenobiotic metabolites out of the cells to be excreted (61). Although expression of xenobiotic metabolizing genes has been previously reported in the endometrium (62), defined and validated endometrial expression and function are still absent. Our data demonstrate transcriptional regulation of genes encoding phase 1 and 2 enzymes, as well as phase 3 transporters in the endometrium. These included numerous aldehyde dehydrogenase (ALDH) members, carboxylesterases, carbohydrate sulfotransferases, cytochrome P450 members, glutathione S-transferases, monoamine oxidases and UDP glycotransferases; and multi-drug resistance protein member ABCC3. Independent qPCR analysis confirmed that xenobiotic metabolism genes were transcriptionally regulated during the menstrual cycle. Interestingly, genes encoding receptors known to mediate xenobiotic metabolism gene expression, including *NR1I3* and *NR1I2* were virtually not expressed (FPKM < 1), while *AHR* was lowly and non-differentially expressed in the endometrium during the phase transition (data not shown), suggesting that transcriptional regulation of the xenobiotic metabolism network may not occur in a classical manner, but rather through alternative regulatory mechanisms (63). Although the impact of xenobiotic metabolism regulation during mid-secretory in the human endometrium remains elusive, there has been evidence linking dysregulation of xenobiotic metabolism genes to pathological conditions such as infertility and cancer (64). Moreover, it has been proposed that xenobiotic metabolism may act as a detoxification mechanism, providing protection and guarding the endometrium against harmful environmental insult for appropriate and efficient implantation, such as environmental estrogen (64).

In addition to xenobiotic metabolism, apoptosis and EMT were also pathways identified as PGR regulated, and both have received ample attention as pathways important in endometrial function. Apoptosis has long been known to mediate uterine homeostasis, a disruption of which is evidently linked to implantation failure and endometriosis (65, 66). Based on our *in silico* analysis, PGR appeared to promote as well as suppress apoptosis in the mid-secretory endometrium (See Supplemental Table 4 (50)). However, the onset of apoptosis in the cycling endometrium is typically around the late-secretory to menstruation phase (67), suggesting a possibility that during the MS phase, PGR acts to balance rather than induce cell death before the mass apoptosis ensues during late-secretory. Indeed, we confirmed increased PGR binding and increased transcription of both pro- and anti-apoptotic genes including *EMP1* (68), *IER3* (69) and *BCL2L10* (70). EMT and its reciprocal pathway, the mesenchymal-epithelial transition (MET) are important modulators of uterine physiology. During each menstrual cycle, the endometrium undergoes extensive remodeling which involves the building and shedding of the functional layer. The origin of the epithelial cells has long been under debate, with some evidence supporting MET being a major player for endometrial re-epithelialization (71, 72). It has been postulated that by retaining imprint of the mesenchymal origin, the endometrial epithelial cells are prone to return to its mesenchymal state via EMT (73). In the MS endometrium, we found various EMT modulating genes to be transcriptionally regulated by PGR, including *MMP2*, *SERPINE1*, *NNMT*, and *WNT5A*. Interestingly, although PGR appeared to promote the expression of EMT genes during the WOI, a closer examination of our gene expression data suggested that the consequences of these regulatory activities resulted in neither decreased epithelial properties nor increased mesenchymal properties. The mesenchymal cell marker *CDH2* was strongly repressed (seven-fold), while another marker, *VIM*, although not identified as a DEG, showed a significant decrease with a fold-change that did not qualify for differential expression in the MS endometrium (data not shown). On the other hand, numerous epithelial cell markers including *CDH1*, *CLDN1*, *CLDN4*, *CLDN8*, *CLDN10*, *KLF4* and *KLF5* were all upregulated during MS. Additionally, *CLDN4*, *CLDN8* and *KLF4* were also presented with increased PGR binding in proximity, suggesting that PGR may directly promote the upregulation of these epithelial markers and maintain the epithelial-like characteristic of these cells. It is possible that while some mesenchymal properties in the epithelium provide for the implanting embryo (such as decreased cell to cell adhesion), but a complete loss of the epithelial status is likely unfavorable and hence PGR acts both to increase EMT as well as maintain the epithelial state. In support of this, EMT has been postulated as an important modulator of noninvasive trophoblast implantation in bovines (74).

### Role and function of the endometrial epithelia during the WOI

RNA-seq was conducted to evaluate the transcriptomic profile in the epithelial compartment of the endometrium. A simple functional annotation found comparable biological functions as that of the whole endometrium, including inflammatory responses, TNFA/NFKB signaling, xenobiotic metabolism, apoptosis, KRAS signaling and EMT as positively enriched; and E2F signaling, G2M checkpoint, mitotic spindle and DNA repair as repressed. IPA predicted the activity of various upstream regulators based on the epithelial transcriptome which included cytokines and transcriptional regulators. Some cytokines identified in our study have been known to facilitate implantation in mammals, whilst the functions of others remain elusive. Additionally, the majority of the epithelial transcription regulators identified in our study have yet to be studied for functional relevance in mediating implantation in the human endometrium, including NUPR1, TBX2, SMARCA4, CEBPA, RABL6 and EP400. Interestingly, the Estrogen Receptors ESR1 and ESR2 showed repression and activation during WOI in the epithelium, respectively. Accordingly, RNA-seq results showed downregulation of ESR1 and upregulation of ESR2 during the phase transition in the epithelium. The repression of ESR1 activity during the window of implantation is well documented, and a mouse model with epithelial ESR1 deletion illustrated a role in regulating apoptosis (75). There is also evidence linking ESR1 overexpression and implantation failure in humans, emphasizing the importance of regulated ESR1 expression during this critical period (76). In contrast, ESR2 has received little attention in the endometrium based on its low expression level compared to the ovary, oviduct or mammary gland (15). Our results showed that there is enrichment of ESR2 expression specifically in the epithelium, with FPKM values in the endometrium averaging 0.75 in the whole endometrium and 2.1 in the epithelium (data not shown). The increased activity of ESR2 (as predicted by IPA), the robust upregulation of its transcript as well as enriched epithelial expression during the WOI propose a possibility that ESR2 engage in previously unrecognized role in mediating pregnancy.

To further unravel molecular pathways with increased specificity to the epithelium, we used two additional approaches. Firstly, we compared the whole endometrial-derived DEGs to epithelial-derived DEGs and excluded the common DEGs to obtain a profile of DEGs that were only detected in the epithelium. Whilst excluding the “common” DEGs may seem counter-intuitive, since the epithelium comprises a part of the endometrium and some “epithelial” genes with substantial transcriptional changes would surface when examined in the whole endometrium, thereby excluding the common DEGs would altogether eliminate those genes. However, the purpose of the epithelial specific examination is to identify previously “missed” epithelial-specific pathways (genes) when examining the endometrium as a whole. Genes in this category may show changes that are subtle but not necessarily less important in nature, and hence our approach of excluding the “common” DEGs. The second approach was to compare the activity status of the upstream regulators calculated for each set of DEGs and identify upstream regulators with enhanced activity in the epithelium. Using the IPA software to examine the 658 epithelial-specific DEGs, the most represented canonical pathways were dermatan sulfate, chondroitin sulfate and cholesterol biosynthesis. Dermatan and chondroitin sulfate are glycosaminoglycans found mostly in the skin, blood vessels and the heart valves (77). They are known to regulate coagulation and wound repair, as well as recruit natural killer cells into the uterus during the reproductive cycle (78). However, the specific role of the endometrial epithelial cells in biosynthesis of these glycosaminoglycans has not yet been reported. On the other hand, progesterone has been reported to inhibit the synthesis of cholesterol in the uterine epithelium of mice, and this has been postulated as a mechanism to block epithelial cell proliferation. Our data accordingly suggest that suppression of cholesterol biosynthesis may be more specifically refined to the epithelial compartment, possibly associated with PGR-mediated inhibition of epithelial cell proliferation during the MS phase (79).

Lastly, we identified two transcription factors, IRF8 (ICSBP) and MEF2C with enhanced activity in the epithelium, whose protein levels and cellular localization were regulated in the epithelium during the WOI. MEF2C specifically showed nuclear localization during this period, and as MEF2C is a transcription factor, it’s likely that the nuclear localization is associated with its increased transcriptional capacity. IRF8 is a member of the interferon (IFN) regulatory factor (IRF) family and is known to regulate gene expression in an interferon-dependent manner (80). It is a modulator of cellular apoptosis under pathological conditions and deregulation of other family members are associated with endometrial adenocarcinoma (81–85), suggesting that IRF proteins may regulate female reproduction. Supporting this, Kashiwagi *et al*. have reported IRF8 expression in the murine endometrium in response to the implanting embryo, but not in pseudopregnancy (86), and Kusama *et al.* later reported the upregulation of IRF8 in the bovine endometrial luminal epithelium in response to the embryo derived interferon tau (87). MEF2C belongs to the MADS box transcription enhancer 2 family, which plays a role in proliferation, invasion and differentiation in various cell types (88). Other members of the family (MEF2A and MEF2D) are known to modulate cytotrophoblast invasion and differentiation in the human placenta (89), and MEF2C itself has been associated with endometriosis, although no apparent function has been reported in the endometrial epithelium (90). Whilst little is known regarding the epithelial function of IRF8 and MEF2C in the endometrium during the WOI, our findings suggest that these factors could have important functions in the uterus and female reproduction.

In summary, signaling pathways controlled by progesterone and PGR are indispensable in uterine biology and homeostasis, a disruption of which manifests in a wide range of gynecological abnormalities such as endometriosis, adenomyosis, fertility defects and endometrial cancer. These pathological conditions are linked to dysregulation of many molecular pathways amongst which are EMT, apoptosis, cell migration and inflammatory response. In this study we provide evidence to show how some of these pathways could be directly controlled by the progesterone signaling through the transcriptional activity of PGR. An understanding of the precise regulatory pattern and mechanism of PGR, that is, what genes are regulated by PGR, and how these genes are regulated by PGR provide a bridging link to explain the molecular mechanism of disease phenotypes under aberrantly regulated PGR conditions. One limitation of this study is that ChIP-seq does not take into consideration the control of PGR over distal DNA response element due to the chromatin interaction in a three-dimensional structure. It is worth noting that roughly 21-22 % of the PGR bound intervals occurred in the “intergenic” regions of the genome (Fig. 1. A), which is defined as greater than 25 kb from the TSS. Although it is possible that these bindings have transcriptional relevance, we cannot draw any conclusion from this study. To address this, future studies should aim to attain a comprehensive three-dimensional structure to elucidate the chromatin conformation in parallel to PGR binding using techniques such as Hi-C (91, 92). This will allow the identification of PGR binding sites in a more global view without the limitation of chromosomal distance. Additional to the PGR regulatory function, approaching the uterine transcriptomic analysis in a compartment specific manner enabled the identification of numerous proteins with previously unrecognized roles in uterine biology and pregnancy. These findings provide a direction for future studies aimed to explore molecular factors crucial for uterine homeostasis.

## Materials and Methods

### Ethics Statement

This project was executed in accordance with the federal regulation governing human subject research. All procedures were approved by the following ethics committees the University of North Carolina at Chapel Hill IRB under file #:05-1757. Informed consent was obtained from all patients before their participation in this study.

### Human Endometrial Samples

We recruited normal volunteers with the following inclusion criteria: ages 18-37, normal menstrual cycle characteristics, an inter-cycle interval of 25-35 days, varying no more than 2 days from cycle to cycle, a normal luteal phase length without luteal spotting, and a body mass index (BMI) between 19 - 28. We excluded women with infertility, pelvic pain, signs and symptoms of endometriosis, history of fibroids or history of taking medication affecting hormonal function in the last 3 months. Endometrial samples were taken using an office biopsy instrument (Pipelle™, Milex Products Inc., Chicago, IL) from the volunteers. Cycle day was determined based on the last menstrual period combined with menstrual history (P samples) or date of Luteinizing Hormone surge. Cycle phase and endometrial normality was confirmed with H&E staining based on the Noyes criteria (93). Details for patients with accessible data are summarized in Supplemental Table 8 (50).

### RNA-seq and Analysis

RNA was prepared from endometrial samples using TRIzol (Thermo Fisher Scientific, Waltham, MA) under the manufacturer’s suggested conditions. Absorption spectroscopy (NanoDrop 8000, Thermo Fisher Scientific, Waltham, MA) was used for quantification of RNA with a ribosomal RNA standard curve. The RNA libraries were sequenced with a HiSeq 2000 system (Illumina). The raw RNA-Seq reads (100 nt, paired-end) were initially processed by filtering with average quality scores greater than 20. Reads which passed the initial processing were aligned to the human reference genome (hg19; Genome Reference Consortium Human Build 19 from February 2009) using TopHat version 2.0.4 (94) and assembled using Cufflinks version 2.0.2 (95). BigWig file was generated from normalized bedgraph file of each sample using bedGraphToBigWig. Scores represent normalized mapped read coverage. Expression values of RNA-Seq were expressed as FPKM (fragments per kilobase of exon per million fragments) values. Differential expression was calculated using Cuffdiff (95). Transcripts with FPKM > 1, *q*-value < 0.05 and at least 1.5-fold change were defined as differentially expressed genes (DEG). The data discussed in this publication have been deposited in NCBI’s Gene Expression Omnibus and are accessible through GEO Series accession number GSE132713 (https://www.ncbi.nlm.nih.gov/geo/query/acc.cgi?acc=GSE132713).

### Chromatin immunoprecipitation sequencing (ChIP-seq) and qRT-PCR (ChIP-qPCR)

Two sets of biopsied tissues were derived from healthy volunteers, each set comprising of one P and one MS endometrial samples (termed P1 and MS1 for set1, and P2 and MS2 for set2). The tissues were flash frozen and sent to the Active Motif company for Factor-Path ChIP-seq analysis. The tissues were fixed, followed by sonication to shear the chromatin into smaller fragments before immunoprecipitation using the Progesterone Receptor (PGR) antibody (sc-7208, Santa Cruz). PGR-bound DNA was subsequently purified and amplified to generate a library for sequencing and quantitative real-time PCR (ChIP-seq and ChIP-qPCR). Sequencing was performed using a NextSeq 500 system (Illumina). The raw ChIP-seq reads (75 nt, single-end) were processed and aligned to the human reference genome (hg19; Genome Reference Consortium Human Build 19 from February 2009) using Bowtie version 1.1.2 (96) with unique mapping and up to 2 mismatches for each read (-m 1 -v 2). The duplicated reads with the same sequence were discarded, and the bigWig files were displayed on UCSC genome browser as custom tracks. Peak calling for each sample was performed by SICER version 1.1 with FDR of 0.001. Software MEDIP was used to identify differential peaks of PGR binding between the P and MS samples (97). Each region was defined as the genomic interval with at least 2-fold difference of read count and *p*-value ≤ 0.01. Each differential peak was mapped to nearby gene using software HOMER’s “annotatePeaks.pl” function (98). As we observed technical variation between sample set1 and set2, we employed a paired-analysis strategy where differential PGR binding intervals were independently determined for P1 VS MS1; and P2 VS MS2. Differential PGR binding that were common to both data sets were used for downstream analysis (Fig. 1. B). Genomic intervals with consistent (or constitutive) PGR binding were defined as motifs bound by PGR in both P and MS phases in either set1, set2 or both; where the read count between the two phases did not qualify for “differential” PGR binding. The motif analysis of differential PGR binding peaks was performed using HOMER software’s “findMotifsGenome.pl” command with default setting (98). The data discussed in this publication have been deposited in NCBI’s Gene Expression Omnibus and are accessible through GEO Series accession number GSE132713 (https://www.ncbi.nlm.nih.gov/geo/query/acc.cgi?acc=GSE132713).

### Epithelial isolation

Endometrial samples obtained from normal controls during the secretory phase of the menstrual cycle were washed with Opti-mem media supplemented with fetal bovine serum (FBS) and antibiotics (10 000 IU/mL penicillin, 10 000 IU/ mL streptomycin; Life Technologies, Grand Island, New York). Tissue was recovered via centrifugation and incubated with collagenase-containing medium (phenol red-free Dul-becco Modified Eagle Medium/F12, 0.5% collagenase I, 0.02% DNase, and 5% FBS). Cell types were separated as described previously (99).

### RNA extraction, cDNA conversion and qPCR

For validation of RNA-seq results, selected genes were examined for RNA expression using independent patients’ samples. Endometrial tissues were resected from patients and flash frozen in liquid nitrogen (see Supplemental Table 8 for patient details (50)). RNA was extracted as described above. Reverse transcription was performed to convert RNA into cDNA using the Moloney Murine Leukemia Virus (MMLV) reverse transcriptase (Thermo Fisher) according to the manufacturer’s instructions. Quantitative real-time PCR was performed using the SsoAdvancedTM Universal SYBR Green Supermix (1725274, Bio-Rad). Briefly, reaction samples were prepared to a total volume of 10 µL with 250 nM of each of the forward and reverse primers, 0.5 ng of cDNA and a final 1 X concentration of the SYBR Green Supermix. The reaction was heated to 98 °C for 30 sec, followed by 35 cycles of denaturation at 95 °C for 5 sec and annealing and elongation for 15 sec. Temperature cycles were performed on the CFX Connect™ Real-Time PCR Detection System (Bio-Rad). The primer sequences were as follows (from 5’ to 3’, F = forward and R = reverse): *CYP3A5* - GTATGAAGGTCAACTCCCTGTG (F) and GGGCCTAAAGACCTTCGATTT (R); *FMO5* - GATTTAAGACCACTGTGTGCAG (F), CCATGACTCCATCAAAGACATTC (R); *UGT1A6* – TGTCTCAGGAATTTGAAGCCTAC (F), GCAATTGCCATAGCTTTCTTCTC (R); *SLCO4A1* – CCCGTCTACATTGCCATCTT (F), GGCCCATTTCCGTGTAGATATT (R); *SLC6A12* – CTTCTACCTGTTCAGCTCCTTC (F), CGTGCAATGCTCTGTGTTC (R); *CYP2C18* – CATTGTGGTGTTGCATGGATATG (F), AGGATTCCAAGTCCTTTGTTAACTT (R); *SULT1C4* – TAAAGCAGGAACAACATGGACT (F), TTCGAGGAAAGGAAATCGTTGA (R); *SLCO2A1* – CTGTACAGCGCCTACTTCAA (F), GATGGCATTGCTGATCTCATTC (R); *GSTM1* – CAAGCACAACCTGTGTGG (F), TTGTCCATGGTCTGGTTCTC (R); *GSTM3* – GGAGTTCACGGATACCTCTTATG (F), GGTAGGGCAGATTAGGAAAGTC (R); *GSTM5* – CGCTTTGAGGGTTTGAAGAAG (F), TGGGCCCTATTTGCTGTT (R); *EMP1* – GTCTTCGTGTTCCAGCTCTT (F), AAGAATGCACAGCCAGCA (R); *IER3* – TGGAACTGCGGCAAAGTA (F), GTAGACAGACGGAGTTGAGATG (R); *BCL2L10* – CCAAAGAACCGCAGAAGAAAC (F), GAAGTTGTGGAGAGATGAGAGG (R); *GPX3* – TCTGGTCATTCTGGGCTTTC (F), ACCTGGTCGGACATACTTGA (R); *TIMP3* – CCCATGTGCAGTACATCCATAC (F), ATCATAGACGCGACCTGTCA (R); *VNN1* – CAGATCAGGGTGCGCATATT (F), GTTTACTTCAGGGTCTGGGATG (R); *SERPINE1* – CTGAGAACTTCAGGATGCAGAT (F), AGACCCTTCACCAAAGACAAG (R); *NNMT* – ACCTCCAAGGACACCTATCT (F), CACACCGTCTAGGCAGAATATC (R); and *TGM2* – ACCCAGCAGGGCTTTATCTA (F), CCCATCTTCAAACTGCCCAA (R). All primers were synthesized by Sigma-Aldrich (St Louis, MO), and gene expression was normalized to 18s rRNA by the ΔΔCT method.

### Immunohistochemistry

Sections were cut from patient’s endometrial biopsies that have been formalin-fixed and paraffin embedded at 5 µm per section. Sections were baked at 65°C for roughly 5 minutes and deparaffined using the Citrisolv clearing agent (22-143-975, Thermo Fisher, Waltham, MA, USA) and hydrated by immersing in decreasing gradient of ethanol. Antigen retrieval was performed using the Vector Labs Antigen Unmasking Solution as per manufacturer’s protocol (H-3300, Vector Laboratories, Burlingame, CA, USA), followed by blocking the endogenous peroxide using 3% hydrogen peroxide diluted in distilled water. Tissues were blocked in 5% normal donkey serum before an overnight incubation with the primary antibody at 4°C (1:200 for ICSBP antibody, sc-365042, Santa Cruz; and 1:100 for MEF2C antibody, SAB4501860, Sigma-Aldrich). The slides were washed twice in PBS at room temperature and applied with secondary antibody diluted 1:200 in 1% BSA prepared in PBS (biotinylated anti-mouse IgG (H+L), BA-9200, and biotinylated anti-rabbit IgG (H+L), BA-1000, Vector Laboratories). The ABC reagent was applied to tissue in accordance with the manufacturer’s instructions (Vector Labs ABC PK-6100, Vector Laboratories). Signal was developed using the Vector Labs DAB ImmPACT staining kit (Vector Labs SK-4105, Vector Laboratories). Finally, the tissue sections were counterstained with hematoxylin and dehydrated through increasing ethanol concentration before incubation in Citrisolv and coverslipping.

### Data Analysis

Various bioinformatic tools were utilized to analyze the high content data generated in this study. Principle component analysis and hierarchical clustering were achieved using the Partek Genomics Suites 7.0 (Partek Inc., St. Louis, MO, USA, http://www.partek.com/partek-genomics-suite/). Functional annotation and enrichment analysis were performed using a combination of the following three tools: Ingenuity Pathway Analysis Software (IPA, http://www.ingenuity.com/), Gene Set Enrichment Analysis (GSEA, http://software.broadinstitute.org/gsea/index.jsp/), and Database for Annotation, Visualization and Integrated Discovery (DAVID, http://david.ncifcrf.gov/). Distribution of PGR binding throughout the genome was conducted using the Peak Annotation and Visualization tool (PAVIS, https://manticore.niehs.nih.gov/pavis2/) (100), and PGR-bound motif was submitted to HOMER motif analysis software to identify presence of other DNA-response elements (http://homer.salk.edu/homer/). GraphPad Prism software was used to analyze single gene expression data generated from both RNA-seq, qPCR, and PGR ChIP-qPCR. Statistical analysis including one-way ANOVA and Student’s *t* test, with a *p-*value of less than 0.05 considered as significant. For pathway analysis using IPA, a given biological category was subjected to Fisher’s exact test to measure the probability that the category was randomly associated. The categories with *p*-values less than 0.05 were defined as significantly enriched.

## Acknowledgements

We thank Dr. Sylvia Hewitt and Dr. John Lydon for editorial assistance. This work was supported by the Intramural Research Program of the National Institute of Health: Project Z1AES103311-01 (F.J.D.), R01HD067721 (S.L.Y.) and 1R01HD096266-01 (T.E.S.).

## Data Availability

All data generated or analyzed during this study are included in this published article or in the data repositories listed in References.

Supplemental tables and figures can be found at: https://doi.org/10.5061/dryad.x69p8czd9

Gene expression data can be found at: https://www.ncbi.nlm.nih.gov/geo/query/acc.cgi?acc=GSE132713

## References

1. Paiva P, Hannan NJ, Hincks C, Meehan KL, Pruysers E, Dimitriadis E, et al. Human chorionic gonadotrophin regulates FGF2 and other cytokines produced by human endometrial epithelial cells, providing a mechanism for enhancing endometrial receptivity. Human reproduction (Oxford, England). 2011;26(5):1153–62.

2. Gellersen B, Brosens JJ. Cyclic decidualization of the human endometrium in reproductive health and failure. Endocrine reviews. 2014;35(6):851–905.

3. Jabbour HN, Kelly RW, Fraser HM, Critchley HO. Endocrine regulation of menstruation. Endocrine reviews. 2006;27(1):17–46.

4. Ramathal CY, Bagchi IC, Taylor RN, Bagchi MK. Endometrial decidualization: of mice and men. Seminars in reproductive medicine. 2010;28(1):17–26.

5. Hawkins SM, Matzuk MM. Menstrual Cycle: Basic Biology. Annals of the New York Academy of Sciences. 2008;1135:10–8.

6. Mesen TB, Young SL. Progesterone and the luteal phase: a requisite to reproduction. Obstetrics and gynecology clinics of North America. 2015;42(1):135–51.

7. Speroff L, Fritz MA. Clinical Gynecologic Endocrinology and Infertility: Lippincott Williams & Wilkins; 2005.

8. Bashiri A, Halper KI, Orvieto R. Recurrent Implantation Failure-update overview on etiology, diagnosis, treatment and future directions. Reproductive biology and endocrinology : RB&E. 2018;16(1):121.

9. Simon A, Laufer N. Assessment and treatment of repeated implantation failure (RIF). Journal of assisted reproduction and genetics. 2012;29(11):1227–39.

10. Garrido-Gomez T, Dominguez F, Quinonero A, Diaz-Gimeno P, Kapidzic M, Gormley M, et al. Defective decidualization during and after severe preeclampsia reveals a possible maternal contribution to the etiology. Proceedings of the National Academy of Sciences of the United States of America. 2017;114(40):E8468–e77.

11. Conrad KP, Rabaglino MB, Post Uiterweer ED. Emerging role for dysregulated decidualization in the genesis of preeclampsia. Placenta. 2017;60:119–29.

12. Macklon NS, Geraedts JP, Fauser BC. Conception to ongoing pregnancy: the ‘black box’ of early pregnancy loss. Human reproduction update. 2002;8(4):333–43.

13. Dey SK, Lim H, Das SK, Reese J, Paria BC, Daikoku T, et al. Molecular cues to implantation. Endocrine reviews. 2004;25(3):341–73.

14. Bhurke AS, Bagchi IC, Bagchi MK. Progesterone-Regulated Endometrial Factors Controlling Implantation. American journal of reproductive immunology (New York, NY : 1989). 2016;75(3):237–45.

15. Pawar S, Hantak AM, Bagchi IC, Bagchi MK. Minireview: Steroid-regulated paracrine mechanisms controlling implantation. Molecular endocrinology (Baltimore, Md). 2014;28(9):1408–22.

16. Young SL. Oestrogen and progesterone action on endometrium: a translational approach to understanding endometrial receptivity. Reproductive biomedicine online. 2013;27(5):497–505.

17. Young SL, Savaris RF, Lessey BA, Sharkey AM, Balthazar U, Zaino RJ, et al. Effect of randomized serum progesterone concentration on secretory endometrial histologic development and gene expression. Human reproduction (Oxford, England). 2017;32(9):1903–14.

18. Wilcox AJ, Baird DD, Weinberg CR. Time of implantation of the conceptus and loss of pregnancy. The New England journal of medicine. 1999;340(23):1796–9.

19. Sunderam S, Kissin DM, Flowers L, Anderson JE, Folger SG, Jamieson DJ, et al. Assisted reproductive technology surveillance--United States, 2009. Morbidity and mortality weekly report Surveillance summaries (Washington, DC : 2002). 2012;61(7):1–23.

20. Patel B, Elguero S, Thakore S, Dahoud W, Bedaiwy M, Mesiano S. Role of nuclear progesterone receptor isoforms in uterine pathophysiology. Human reproduction update. 2015;21(2):155–73.

21. Yin P, Roqueiro D, Huang L, Owen JK, Xie A, Navarro A, et al. Genome-wide progesterone receptor binding: cell type-specific and shared mechanisms in T47D breast cancer cells and primary leiomyoma cells. PloS one. 2012;7(1):e29021.

22. Kim JJ, Kurita T, Bulun SE. Progesterone action in endometrial cancer, endometriosis, uterine fibroids, and breast cancer. Endocrine reviews. 2013;34(1):130–62.

23. Tamm K, Rõõm M, Salumets A, Metsis M. Genes targeted by the estrogen and progesterone receptors in the human endometrial cell lines HEC1A and RL95-2. Reproductive biology and endocrinology : RB&E. 2009;7:150.

24. Matsumoto H, Zhao X, Das SK, Hogan BL, Dey SK. Indian hedgehog as a progesterone-responsive factor mediating epithelial-mesenchymal interactions in the mouse uterus. Developmental biology. 2002;245(2):280–90.

25. Takamoto N, Zhao B, Tsai SY, DeMayo FJ. Identification of Indian hedgehog as a progesterone-responsive gene in the murine uterus. Molecular endocrinology (Baltimore, Md). 2002;16(10):2338–48.

26. Franco HL, Rubel CA, Large MJ, Wetendorf M, Fernandez-Valdivia R, Jeong JW, et al. Epithelial progesterone receptor exhibits pleiotropic roles in uterine development and function. FASEB journal : official publication of the Federation of American Societies for Experimental Biology. 2012;26(3):1218–27.

27. Pan H, Zhu L, Deng Y, Pollard JW. Microarray analysis of uterine epithelial gene expression during the implantation window in the mouse. Endocrinology. 2006;147(10):4904–16.

28. Mazur EC, Vasquez YM, Li X, Kommagani R, Jiang L, Chen R, et al. Progesterone receptor transcriptome and cistrome in decidualized human endometrial stromal cells. Endocrinology. 2015;156(6):2239–53.

29. Li Q, Kannan A, DeMayo FJ, Lydon JP, Cooke PS, Yamagishi H, et al. The antiproliferative action of progesterone in uterine epithelium is mediated by Hand2. Science (New York, NY). 2011;331(6019):912–6.

30. Lee KY, Jeong JW, Wang J, Ma L, Martin JF, Tsai SY, et al. Bmp2 is critical for the murine uterine decidual response. Molecular and cellular biology. 2007;27(15):5468–78.

31. Li Q, Kannan A, Wang W, Demayo FJ, Taylor RN, Bagchi MK, et al. Bone morphogenetic protein 2 functions via a conserved signaling pathway involving Wnt4 to regulate uterine decidualization in the mouse and the human. The Journal of biological chemistry. 2007;282(43):31725–32.

32. Lim H, Ma L, Ma WG, Maas RL, Dey SK. Hoxa-10 regulates uterine stromal cell responsiveness to progesterone during implantation and decidualization in the mouse. Molecular endocrinology (Baltimore, Md). 1999;13(6):1005–17.

33. Taylor HS, Arici A, Olive D, Igarashi P. HOXA10 is expressed in response to sex steroids at the time of implantation in the human endometrium. The Journal of clinical investigation. 1998;101(7):1379–84.

34. Kannan A, Fazleabas AT, Bagchi IC, Bagchi MK. The transcription factor C/EBPbeta is a marker of uterine receptivity and expressed at the implantation site in the primate. Reproductive sciences (Thousand Oaks, Calif). 2010;17(5):434–43.

35. Mantena SR, Kannan A, Cheon YP, Li Q, Johnson PF, Bagchi IC, et al. C/EBPbeta is a critical mediator of steroid hormone-regulated cell proliferation and differentiation in the uterine epithelium and stroma. Proceedings of the National Academy of Sciences of the United States of America. 2006;103(6):1870–5.

36. Plante BJ, Kannan A, Bagchi MK, Yuan L, Young SL. Cyclic regulation of transcription factor C/EBP beta in human endometrium. Reproductive biology and endocrinology : RB&E. 2009;7:15.

37. Gray CA, Bartol FF, Tarleton BJ, Wiley AA, Johnson GA, Bazer FW, et al. Developmental biology of uterine glands. Biology of reproduction. 2001;65(5):1311–23.

38. Kelleher AM, DeMayo FJ, Spencer TE. Uterine Glands: Developmental Biology and Functional Roles in Pregnancy. Endocrine reviews. 2019.

39. Singh H, Aplin JD. Adhesion molecules in endometrial epithelium: tissue integrity and embryo implantation. Journal of anatomy. 2009;215(1):3–13.

40. Wang B, Ye TM, Lee KF, Chiu PC, Pang RT, Ng EH, et al. Annexin A2 Acts as an Adhesion Molecule on the Endometrial Epithelium during Implantation in Mice. PloS one. 2015;10(10):e0139506.

41. Guffanti E, Kittur N, Brodt ZN, Polotsky AJ, Kuokkanen SM, Heller DS, et al. Nuclear pore complex proteins mark the implantation window in human endometrium. Journal of cell science. 2008;121(Pt 12):2037–45.

42. Salker MS, Steel JH, Hosseinzadeh Z, Nautiyal J, Webster Z, Singh Y, et al. Activation of SGK1 in Endometrial Epithelial Cells in Response to PI3K/AKT Inhibition Impairs Embryo Implantation. Cellular physiology and biochemistry : international journal of experimental cellular physiology, biochemistry, and pharmacology. 2016;39(5):2077–87.

43. Boeddeker SJ, Hess AP. The role of apoptosis in human embryo implantation. Journal of reproductive immunology. 2015;108:114–22.

44. Joswig A, Gabriel HD, Kibschull M, Winterhager E. Apoptosis in uterine epithelium and decidua in response to implantation: evidence for two different pathways. Reproductive biology and endocrinology : RB&E. 2003;1:44.

45. Valdez-Morales FJ, Gamboa-Domínguez A, Vital-Reyes VS, Cruz JC, Chimal-Monroy J, Franco-Murillo Y, et al. Changes in receptivity epithelial cell markers of endometrium after ovarian stimulation treatments: its role during implantation window. Reproductive health. 2015;12:45.

46. Kelleher AM, Milano-Foster J, Behura SK, Spencer TE. Uterine glands coordinate on-time embryo implantation and impact endometrial decidualization for pregnancy success. Nature communications. 2018;9(1):2435.

47. Aghajanova L. Leukemia inhibitory factor and human embryo implantation. Annals of the New York Academy of Sciences. 2004;1034:176–83.

48. Kimber SJ. Leukaemia inhibitory factor in implantation and uterine biology. Reproduction (Cambridge, England). 2005;130(2):131–45.

49. Norwitz ER, Schust DJ, Fisher SJ. Implantation and the survival of early pregnancy. The New England journal of medicine. 2001;345(19):1400–8.

50. Chi RA, Wang T, Adams NR, Wu S, Young SL, Spencer TE, et al. Human Endometrial Transcriptome and Progesterone Receptor Cistrome Reveal Important Pathways and Epithelial Regulators. Dryad. 2019;Dataset(https://doi.org/10.5061/dryad.x69p8czd9).

51. Vassen L, Wegrzyn W, Klein-Hitpass L. Human insulin receptor substrate-2 (IRS-2) is a primary progesterone response gene. Molecular endocrinology (Baltimore, Md). 1999;13(3):485–94.

52. Kaya HS, Hantak AM, Stubbs LJ, Taylor RN, Bagchi IC, Bagchi MK. Roles of progesterone receptor A and B isoforms during human endometrial decidualization. Molecular endocrinology (Baltimore, Md). 2015;29(6):882–95.

53. Okada H, Tsuzuki T, Murata H. Decidualization of the human endometrium. Reproductive medicine and biology. 2018;17(3):220–7.

54. Rubel CA, Wu SP, Lin L, Wang T, Lanz RB, Li X, et al. A Gata2-Dependent Transcription Network Regulates Uterine Progesterone Responsiveness and Endometrial Function. Cell reports. 2016;17(5):1414–25.

55. Wang X, Li X, Wang T, Wu SP, Jeong JW, Kim TH, et al. SOX17 regulates uterine epithelial-stromal cross-talk acting via a distal enhancer upstream of Ihh. Nature communications. 2018;9(1):4421.

56. Huang da W, Sherman BT, Lempicki RA. Bioinformatics enrichment tools: paths toward the comprehensive functional analysis of large gene lists. Nucleic acids research. 2009;37(1):1–13.

57. Huang da W, Sherman BT, Lempicki RA. Systematic and integrative analysis of large gene lists using DAVID bioinformatics resources. Nature protocols. 2009;4(1):44–57.

58. Subramanian A, Tamayo P, Mootha VK, Mukherjee S, Ebert BL, Gillette MA, et al. Gene set enrichment analysis: a knowledge-based approach for interpreting genome-wide expression profiles. Proceedings of the National Academy of Sciences of the United States of America. 2005;102(43):15545–50.

59. Owen GI, Richer JK, Tung L, Takimoto G, Horwitz KB. Progesterone regulates transcription of the p21(WAF1) cyclin-dependent kinase inhibitor gene through Sp1 and CBP/p300. The Journal of biological chemistry. 1998;273(17):10696–701.

60. Tseng L, Tang M, Wang Z, Mazella J. Progesterone receptor (hPR) upregulates the fibronectin promoter activity in human decidual fibroblasts. DNA and cell biology. 2003;22(10):633–40.

61. Omiecinski CJ, Vanden Heuvel JP, Perdew GH, Peters JM. Xenobiotic metabolism, disposition, and regulation by receptors: from biochemical phenomenon to predictors of major toxicities. Toxicological sciences : an official journal of the Society of Toxicology. 2011;120 Suppl 1:S49–75.

62. Tseng LH, Chen I, Chen MY, Yan H, Wang CN, Lee CL. Genome-based expression profiling as a single standardized microarray platform for the diagnosis of endometrial disorder: an array of 126-gene model. Fertility and sterility. 2010;94(1):114–9.

63. Hariparsad N, Chu X, Yabut J, Labhart P, Hartley DP, Dai X, et al. Identification of pregnane-X receptor target genes and coactivator and corepressor binding to promoter elements in human hepatocytes. Nucleic acids research. 2009;37(4):1160–73.

64. Altmae S, Martinez-Conejero JA, Salumets A, Simon C, Horcajadas JA, Stavreus-Evers A. Endometrial gene expression analysis at the time of embryo implantation in women with unexplained infertility. Molecular human reproduction. 2010;16(3):178–87.

65. Harada T, Kaponis A, Iwabe T, Taniguchi F, Makrydimas G, Sofikitis N, et al. Apoptosis in human endometrium and endometriosis. Human reproduction update. 2004;10(1):29–38.

66. Antsiferova Y, Sotnikova N. Apoptosis and endometrial receptivity: Relationship with in vitro fertilization treatment outcome. World Journal of Obstetrics and Gynecology. 2016;5(1):87–96.

67. Kokawa K, Shikone T, Nakano R. Apoptosis in the human uterine endometrium during the menstrual cycle. The Journal of clinical endocrinology and metabolism. 1996;81(11):4144–7.

68. Li H, Zhang X, Jiang X, Ji X. The expression and function of epithelial membrane protein 1 in laryngeal carcinoma. International journal of oncology. 2017;50(1):141–8.

69. Akilov OE, Wu MX, Ustyugova IV, Falo LD, Jr., Geskin LJ. Resistance of Sezary cells to TNF-alpha-induced apoptosis is mediated in part by a loss of TNFR1 and a high level of the IER3 expression. Experimental dermatology. 2012;21(4):287–92.

70. Opferman JT, Kothari A. Anti-apoptotic BCL-2 family members in development. Cell death and differentiation. 2018;25(1):37–45.

71. Maybin JA, Critchley HO. Menstrual physiology: implications for endometrial pathology and beyond. Human reproduction update. 2015;21(6):748–61.

72. Bilyk O, Coatham M, Jewer M, Postovit LM. Epithelial-to-Mesenchymal Transition in the Female Reproductive Tract: From Normal Functioning to Disease Pathology. Frontiers in oncology. 2017;7:145.

73. Matsuzaki S, Darcha C. Epithelial to mesenchymal transition-like and mesenchymal to epithelial transition-like processes might be involved in the pathogenesis of pelvic endometriosis. Human reproduction (Oxford, England). 2012;27(3):712–21.

74. Bai R, Kusama K, Nakamura K, Sakurai T, Kimura K, Ideta A, et al. Down-regulation of transcription factor OVOL2 contributes to epithelial-mesenchymal transition in a noninvasive type of trophoblast implantation to the maternal endometrium. FASEB journal : official publication of the Federation of American Societies for Experimental Biology. 2018:fj201701131RR.

75. Winuthayanon W, Hewitt SC, Orvis GD, Behringer RR, Korach KS. Uterine epithelial estrogen receptor alpha is dispensable for proliferation but essential for complete biological and biochemical responses. Proceedings of the National Academy of Sciences of the United States of America. 2010;107(45):19272–7.

76. Dorostghoal M, Ghaffari HO, Marmazi F, Keikhah N. Overexpression of Endometrial Estrogen Receptor-Alpha in The Window of Implantation in Women with Unexplained Infertility. International journal of fertility & sterility. 2018;12(1):37–42.

77. Trowbridge JM, Gallo RL. Dermatan sulfate: new functions from an old glycosaminoglycan. Glycobiology. 2002;12(9):117r–25r.

78. Kitaya K, Yasuo T. Regulatory role of membrane-bound form interleukin-15 on human uterine microvascular endothelial cells in circulating CD16(-) natural killer cell extravasation into human endometrium. Biology of reproduction. 2013;89(3):70.

79. Stacey K, Beasley B, Wilce PA, Martin L. Effects of female sex hormones on lipid metabolism in the uterine epithelium of the mouse. The International journal of biochemistry. 1991;23(3):371–6.

80. Tailor P, Tamura T, Kong HJ, Kubota T, Kubota M, Borghi P, et al. The feedback phase of type I interferon induction in dendritic cells requires interferon regulatory factor 8. Immunity. 2007;27(2):228–39.

81. Hu X, Yang D, Zimmerman M, Liu F, Yang J, Kannan S, et al. IRF8 regulates acid ceramidase expression to mediate apoptosis and suppresses myelogeneous leukemia. Cancer research. 2011;71(8):2882–91.

82. Liu K, Abrams SI. Coordinate regulation of IFN consensus sequence-binding protein and caspase-1 in the sensitization of human colon carcinoma cells to Fas-mediated apoptosis by IFN-gamma. Journal of immunology (Baltimore, Md : 1950). 2003;170(12):6329–37.

83. Yang D, Thangaraju M, Greeneltch K, Browning DD, Schoenlein PV, Tamura T, et al. Repression of IFN regulatory factor 8 by DNA methylation is a molecular determinant of apoptotic resistance and metastatic phenotype in metastatic tumor cells. Cancer research. 2007;67(7):3301–9.

84. Giatromanolaki A, Koukourakis MI, Ritis K, Mimidis K, Sivridis E. Interferon regulatory factor-1 (IRF-1) suppression and derepression during endometrial tumorigenesis and cancer progression. Cytokine. 2004;26(4):164–8.

85. Kuroboshi H, Okubo T, Kitaya K, Nakayama T, Daikoku N, Fushiki S, et al. Interferon regulatory factor-1 expression in human uterine endometrial carcinoma. Gynecologic oncology. 2003;91(2):354–8.

86. Kashiwagi A, DiGirolamo CM, Kanda Y, Niikura Y, Esmon CT, Hansen TR, et al. The Postimplantation Embryo Differentially Regulates Endometrial Gene Expression and Decidualization. Endocrinology. 2007;148(9):4173–84.

87. Kusama K, Bai R, Nakamura K, Okada S, Yasuda J, Imakawa K. Endometrial factors similarly induced by IFNT2 and IFNTc1 through transcription factor FOXS1. PloS one. 2017;12(2):e0171858.

88. Pon JR, Marra MA. MEF2 transcription factors: developmental regulators and emerging cancer genes. Oncotarget. 2016;7(3):2297–312.

89. Li L, Rubin LP, Gong X. MEF2 transcription factors in human placenta and involvement in cytotrophoblast invasion and differentiation. Physiological genomics. 2018;50(1):10–9.

90. Ohlsson Teague EM, Van der Hoek KH, Van der Hoek MB, Perry N, Wagaarachchi P, Robertson SA, et al. MicroRNA-regulated pathways associated with endometriosis. Molecular endocrinology (Baltimore, Md). 2009;23(2):265–75.

91. de Wit E, de Laat W. A decade of 3C technologies: insights into nuclear organization. Genes & development. 2012;26(1):11–24.

92. Diaz N, Kruse K, Erdmann T, Staiger AM, Ott G, Lenz G, et al. Chromatin conformation analysis of primary patient tissue using a low input Hi-C method. Nature communications. 2018;9(1):4938.

93. Noyes RW, Hertig AT, Rock J. Dating the endometrial biopsy. American journal of obstetrics and gynecology. 1975;122(2):262–3.

94. Trapnell C, Pachter L, Salzberg SL. TopHat: discovering splice junctions with RNA-Seq. Bioinformatics (Oxford, England). 2009;25(9):1105–11.

95. Trapnell C, Williams BA, Pertea G, Mortazavi A, Kwan G, van Baren MJ, et al. Transcript assembly and quantification by RNA-Seq reveals unannotated transcripts and isoform switching during cell differentiation. Nature biotechnology. 2010;28(5):511–5.

96. Langmead B, Trapnell C, Pop M, Salzberg SL. Ultrafast and memory-efficient alignment of short DNA sequences to the human genome. Genome biology. 2009;10(3):R25.

97. Zang C, Schones DE, Zeng C, Cui K, Zhao K, Peng W. A clustering approach for identification of enriched domains from histone modification ChIP-Seq data. Bioinformatics (Oxford, England). 2009;25(15):1952–8.

98. Heinz S, Benner C, Spann N, Bertolino E, Lin YC, Laslo P, et al. Simple combinations of lineage-determining transcription factors prime cis-regulatory elements required for macrophage and B cell identities. Molecular cell. 2010;38(4):576–89.

99. Arnold JT, Kaufman DG, Seppala M, Lessey BA. Endometrial stromal cells regulate epithelial cell growth in vitro: a new co-culture model. Human reproduction (Oxford, England). 2001;16(5):836–45.

100. Huang W, Loganantharaj R, Schroeder B, Fargo D, Li L. PAVIS: a tool for Peak Annotation and Visualization. Bioinformatics (Oxford, England). 2013;29(23):3097–9.

