## Supplementary figures and images for "Human Endometrial Transcriptome and Progesterone Receptor Cistrome Reveal Important Pathways and Epithelial Regulators"

### Supplemental Figures

#### Slide 1
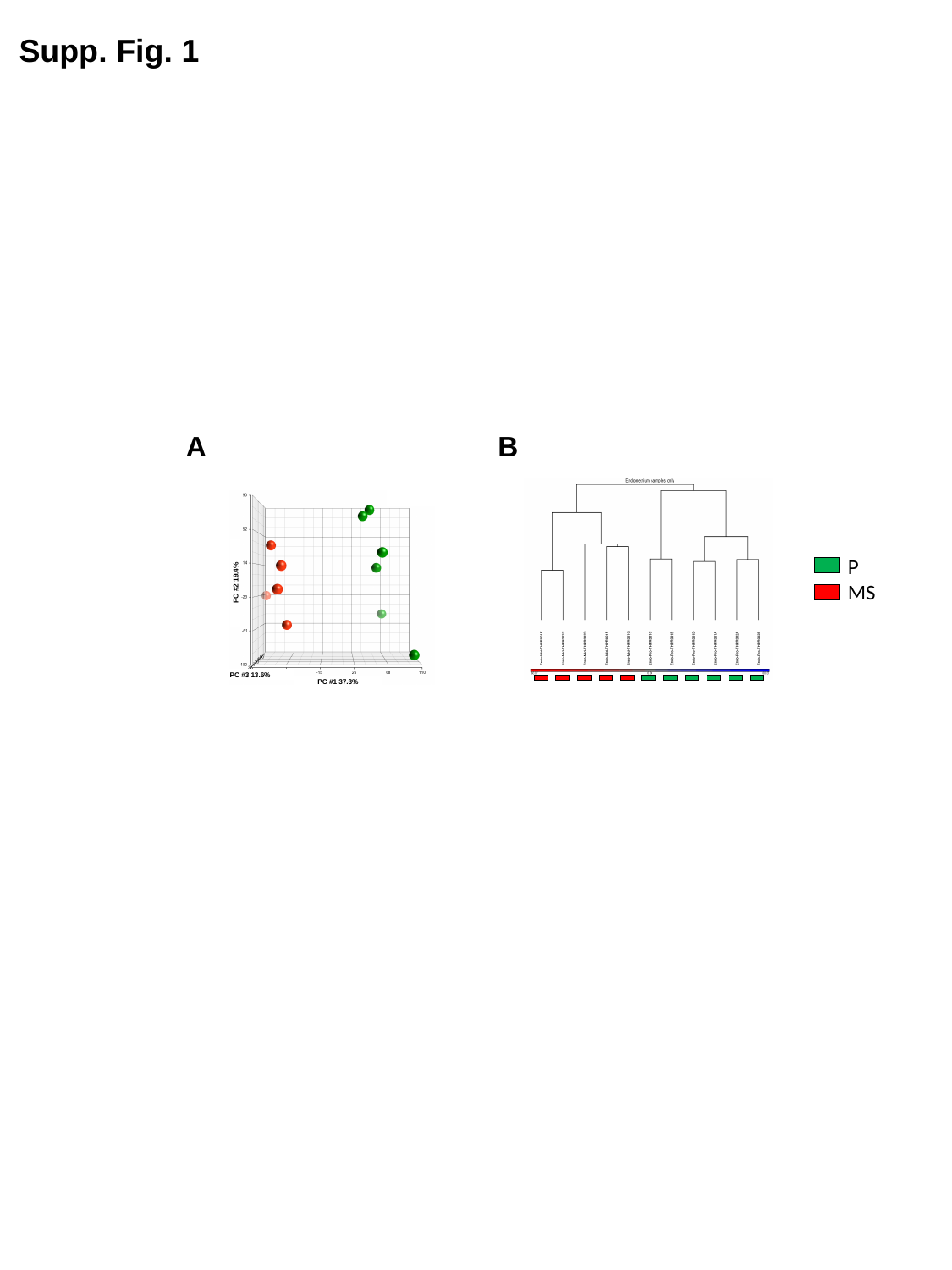

Supp. Fig. 1
B
A
P
MS
PC #2 19.4%
PC #3 13.6%
PC #1 37.3%

#### Slide 2
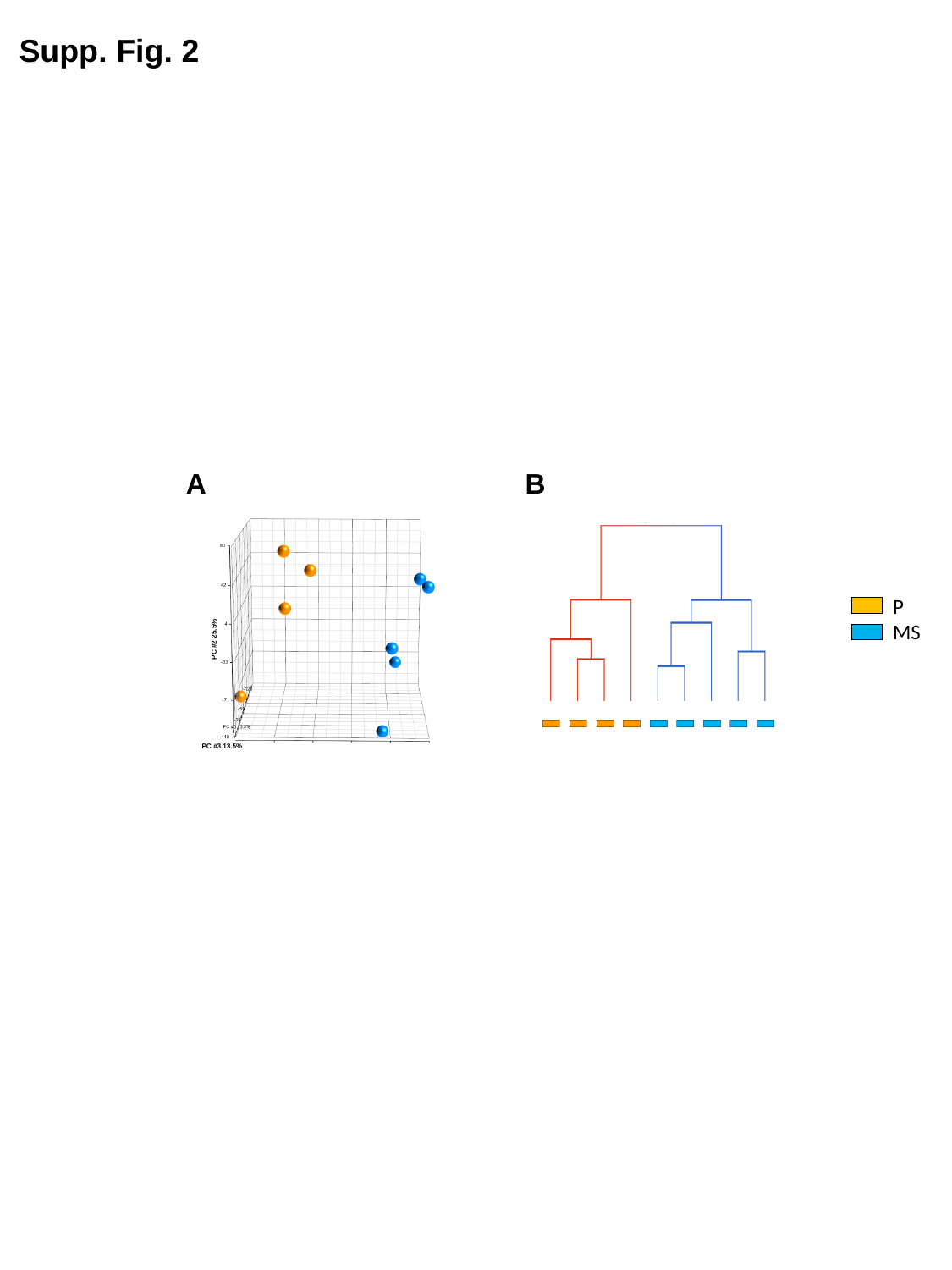

Supp. Fig. 2
B
A
PC #2 25.5%
PC #3 13.5%
P
MS
